# Dynamic evolution of diaphorin biosynthetic genes against a background of genome conservation in *Profftella*

**DOI:** 10.64898/2026.10.30.762998

**Authors:** Yuka Yasuda, Jörn Piel, Igor Malenovský, Ilia Gjonov, Yuu Hirose, Atsushi Nakabachi

## Abstract

The Asian citrus psyllid *Diaphorina citri* (Insecta: Hemiptera), a major agricultural pest, harbors the unusual obligate bacterial symbiont *“Candidatus* Profftella armatura” (*Profftella*), which is intimately integrated with its host yet devotes a remarkably large proportion of its highly reduced genome to the production of diaphorin, a bioactive polyketide. To gain insight into the evolution of this distinctive symbiont, we determined the genomes of *Profftella* and the co-resident nutritional symbiont *“Candidatus* Carsonella ruddii” (*Carsonella*) from *Diaphorina lycii* and compared them with previously characterized counterparts from *D. citri* and *D*. cf. *continua*. The *Profftella* and *Carsonella* chromosomes from *D. lycii* were 471 and 174 kb, respectively, and *Profftella* harbored an 8-kb plasmid. Genome organization and gene content were highly conserved across the three lineages. The diaphorin biosynthetic system likewise retained its overall organization, consistent with an important role for diaphorin in the *Diaphorina* symbiotic system. Remarkably, however, pronounced lineage-specific structural variation was observed in diaphorin synthases DipP and DipT, encoded by two of the 20 diaphorin biosynthetic genes. These synthases contained three extensively amplified tandem arrays of acyl carrier protein (ACP) domains that carry growing polyketide intermediates, with up to seven copies compared with one or two at corresponding positions in the related pederin synthases. All three regions of tandem ACP repeats showed copy-number variation among the *Profftella* lineages. This variation indicates that ACP amplification remains evolutionarily dynamic despite the otherwise highly constrained symbiotic genome. Together, our findings show that even highly integrated obligate symbionts retain evolutionary flexibility in specialized functional systems.

## Introduction

Bacterial symbionts of animals provide their hosts with diverse benefits, including nutritional supplementation and defense against natural enemies [1, 2]. In insects, nutritional symbionts are particularly important for hosts subsisting on nutritionally deficient diets and often form long-term obligate associations with their hosts. Such symbionts are vertically transmitted, with many residing in specialized organs called bacteriomes. These host-dependent associations are often accompanied by drastic genome reduction, sometimes to well below 1 Mb, while genes for the biosynthesis of host-essential nutrients are retained [2, 3]. Defensive symbionts, by contrast, protect their hosts against predators, parasitoids, pathogens, or other natural enemies through bioactive compounds, but are generally less stably associated with their hosts than nutritional symbionts and rarely show the extreme genome reduction associated with long-term symbiosis [4, 5].

Psyllids (Hemiptera: Psylloidea) are plant sap-feeding insects that harbor vertically transmitted bacterial symbionts in their bacteriomes [6, 7]. They universally harbor the obligate nutritional symbiont “*Candidatus* Carsonella ruddii” (hereafter *Carsonella*; Gammaproteobacteria) as their primary symbiont. Phylogenetic evidence indicates that *Carsonella* was acquired by a common ancestor of psyllids and has since undergone cospeciation with its hosts [8, 9]. *Carsonella* has one of the smallest known cellular genomes and supplies its host with essential amino acids deficient in the phloem-sap diet [10–20]. Many psyllid lineages additionally harbor another obligate symbiont, collectively referred to as secondary symbionts, which have been acquired independently from diverse bacterial lineages during psyllid evolution [11–18, 21–23]. The Asian citrus psyllid, *Diaphorina citri* (Psyllidae), an important agricultural pest and vector of “*Candidatus* Liberibacter” spp. (Alphaproteobacteria) [24–26], harbors “*Candidatus* Profftella armatura” (hereafter *Profftella*; Betaproteobacteria) as its secondary symbiont [12, 27–30]. Like typical obligate nutritional symbionts, *Profftella* has an extremely reduced genome of approximately 460 kb, yet retains little apparent capacity for host nutritional provisioning: its remaining nutrition-related functions are largely limited to riboflavin, biotin, and carotenoid biosynthesis, functions that are often dispensable and lost in many other symbionts of plant sap-feeding insects. Most notably, a large fraction of its chromosomal coding capacity is instead devoted to the biosynthesis of diaphorin, a bioactive polyketide of the pederin family [12]. Diaphorin is toxic to various organisms, including natural enemies of *D. citri*, strongly implicating it in host defense [12, 31–37]. Thus, *Profftella* represents an unusual obligate symbiont in which an exceptionally intimate association with its host, reflected in bacteriome localization, strict vertical transmission, and extreme genome reduction, is coupled with a prominent role in host defense through specialized secondary metabolism.

16S rRNA gene amplicon analyses of psyllid bacterial communities subsequently identified closely related *Profftella* lineages in two other *Diaphorina* species, *D*. cf. *continua* and *D. lycii* [38]. Comparison of the complete *Profftella* and *Carsonella* genomes from *D*. cf. *continua* with their counterparts from *D. citri* revealed extensive conservation of genome organization and gene content [13]. The diaphorin biosynthetic system, comprising 20 genes (*dipA*–*T*) in two chromosomal clusters, was likewise highly conserved. A distinctive feature of this system was the extensive tandem amplification of acyl carrier protein (ACP) domains in DipP and DipT, two large multidomain polyketide synthases [13].

Although closely related, the three *Diaphorina* species differ markedly in their ecology and geographic distributions. *D. citri* is native to tropical and subtropical South to East Asia and feeds on rutaceous plants including *Citrus* spp. [24], whereas *D*. cf. *continua* occurs in the Mediterranean region and is associated with silvery spurge flax (*Thymelaea tartonraira*, Thymelaeaceae) [38]. *D. lycii* has a broad distribution from Southern Europe and North Africa through the Middle East and Central Asia to Mongolia and is narrowly oligophagous on *Lycium* spp. (Solanaceae) [39]. Here, we determined the complete genomes of *Profftella* and *Carsonella* from *D. lycii* and compared them with their previously characterized counterparts from *D. citri* and *D*. cf. *continua*. Hereafter, the suffixes _Dly, _DC, and _Dco denote symbiont lineages associated with *D. lycii*, *D. citri*, and *D*. cf. *continua*, respectively (e.g., *Profftella*_Dly and *Carsonella*_Dly).

## Materials and Methods

### Insects and DNA preparation

Adult *D. lycii* were collected from *Lycium barbarum* (Solanaceae) in the floodplain of the Stara River, 2.3 km northeast of Byaga village, Pazardzhik Province, Bulgaria (42°4′45″N, 24°24′11″E; 250 m a.s.l.), on June 18, 2017. Bacteriomes were dissected, and genomic DNA was extracted using the DNeasy Blood & Tissue Kit (Qiagen). DNA quality was assessed using a NanoDrop 2000c spectrophotometer (Thermo Fisher Scientific), and DNA concentration was quantified using a Qubit 2.0 Fluorometer with the Qubit dsDNA HS Assay Kit (Thermo Fisher Scientific).

### Genome sequencing, assembly, and finishing

A paired-end sequencing library was prepared from genomic DNA using the KAPA HyperPlus Kit (Roche), with enzymatic fragmentation performed at 37°C for 40 min and adapter ligation extended to 30 min; all other procedures followed the manufacturer’s protocol. The library was sequenced on an Illumina HiSeq X platform with 150-bp paired-end reads. Genome assemblies were generated using SPAdes v4.1.0 [40] and Unicycler v0.5.1 [41] and inspected with Bandage v0.9.0 [42]. Ambiguous regions were resolved by Sanger sequencing.

### Genome annotation and comparative genomic analysis

Gene prediction and functional annotation were performed using DFAST v1.2.0 [43] and PGAP 2023-05-17.build6771 [44]. Genes remaining uncharacterized after automated annotation were examined by BLAST+ v2.17.0 searches [45]. NCBI ORFfinder [46] was used to identify genes missed by the automated annotation and to refine gene boundaries where appropriate. COG categories were assigned using eggNOG-mapper v3.0.0 [47] with orthology assignments from the eggNOG 7 database [48]. To investigate functions potentially beneficial to the host psyllids, the genomes were queried against KEGG (Release 118) [49].

Comparative analyses included the newly determined *Carsonella*_Dly and *Profftella*_Dly genomes, together with the previously reported genomes from *D. citri* (*Profftella*_DC, GCA_000441555.1; *Carsonella*_DC, GCA_000441575.1) and *D*. cf. *continua* (*Profftella*_Dco, GCA_016593155.1; *Carsonella*_Dco, GCA_016593055.1). Genome-wide nucleotide similarities and synteny were examined by all-against-all BLASTN searches in GenomeMatcher v3.06 [50] using the parameters “-F F -W 21 -e 1.0e-10”. Average nucleotide identity (ANI) was calculated with the OrthoANIu algorithm implemented in the ANI Calculator (EzBioCloud) [51], using reciprocal best hits, a 1,000-bp window, and a 200-bp step.

### Phylogenetic analyses

A whole-genome-based phylogenetic analysis was performed for 17 bacterial lineages, including *Carsonella* and *Profftella* from the three *Diaphorina* species. Single-copy orthologs were identified with OrthoFinder v2.5.5 [52] and aligned using MAFFT v7.526 [53]. A concatenated supermatrix was constructed with catsequences v1.5. The LG+I+G4+F model was selected using ModelTest-NG v0.1.7 [54], and maximum-likelihood inference was performed with RAxML-NG v1.2.1 [55] using 1,000 bootstrap replicates. Trees were visualized with iTOL v7.2.2 [56]. Additional maximum-likelihood phylogenetic analyses were performed for phytoene desaturase (CrtI) and ketoacyl synthase (KS) domains using MAFFT v7.526 and RAxML-NG v1.2.1, with substitution models selected by ModelTest-NG v0.1.7.

### Comparison of metabolic gene repertoires

Genes involved in essential amino-acid, sulfur, biotin, and riboflavin metabolism were compared among the *Diaphorina*-associated *Carsonella* and *Profftella* lineages and, for broader comparison, *Carsonella* and secondary symbionts from *Anomoneura mori* and *Cacopsylla melanoneura* (Psyllidae). The secondary symbiont of *C. melanoneura* is “*Candidatus* Psyllophila symbiotica” (hereafter *Psyllophila*; Gammaproteobacteria) [14, 16], which is widespread among *Cacopsylla* species and closely related to the secondary symbiont of *A. mori* [15, 57]. Published data from additional *Psyllophila* lineages were also considered when assessing the conservation of nutrient-biosynthetic functions and sulfur-metabolism genes [14]. Presence/absence patterns were evaluated from the genome annotations and pathway assignments described above and from the cited studies.

### Substitution-rate and *dN*/*dS* analyses

Orthologous CDSs shared among the symbiont genomes from *D. lycii*, *D. citri*, and *D*. cf. *continua* were translated and aligned with MAFFT v7.526 using the E-INS-i algorithm with default parameters. Protein alignments were converted to codon-based nucleotide alignments using PAL2NAL v14 [58]. Nonsynonymous (*dN*) and synonymous (*dS*) substitution rates for orthologous pairs were calculated with KaKs_Calculator 2.0 [59] using the YN model, and *dN*/*dS* ratios were calculated from these estimates. Statistical analyses and plotting were performed in R v4.5.1 [60] using the brunnermunzel, ggplot2, and ggExtra packages. For each of the Dly–DC and Dly–Dco lineage pairs, distributions of *dN* and *dS* were compared between *Carsonella* and *Profftella* using the Brunner–Munzel test. Genome-wide distributions of *dN*/*dS* ratios were also examined for each symbiont.

### Analysis of diaphorin biosynthetic enzymes

Domain architectures of the diaphorin synthases were analyzed with TransATor [61] and compared among the three *Profftella* lineages and with homologous pederin synthases from the *Pseudomonas* endosymbiont of *Paederus fuscipes*.

## Results

### Genome-wide synteny is strongly conserved in both *D. lycii* symbionts

Illumina short-read sequencing provided mean coverages of 344.4× for *Profftella*_Dly and 78.2× for *Carsonella*_Dly. Sanger sequencing resolved remaining ambiguities, yielding complete genomes for both symbionts. The *Profftella*_Dly chromosome was 471,402 bp (25.2% G+C) and its circular plasmid was 8,128 bp (25.0% G+C). The chromosome encoded 371 predicted CDSs, one rRNA operon, and 34 tRNAs, while the plasmid encoded four CDSs, three of which were involved in carotenoid biosynthesis. The *Carsonella*_Dly chromosome was 173,860 bp (17.8% G+C), with 205 predicted CDSs, one rRNA operon, and 27 tRNAs (Table 1; Fig. S1; Tables S1 and S2).

**Table 1.** Genomic features of *Carsonella* and *Profftella* from three *Diaphorina* species.

|  | ***Profftella*_Dly** | ***Profftella*_DC** | ***Profftella*_Dco** | ***Carsonella*_Dly** | ***Carsonella*_DC** | ***Carsonella*_Dco** |
|---|---|---|---|---|---|---|
| Chromosome size (bp) | 471,402 | 459,399 | 469,264 | 173,860 | 174,014 | 173,853 |
| Plasmid size (bp) | 8,128 | 5,458 | >5,952 | - | - | - |
| G+C content (%) | 25.2 [25.0] | 24.2 [23.9] | 24.4 [23.1] | 17.8 | 17.6 | 17.9 |
| CDSs | 371 [4] | 365 [6] | 355 [4] | 205 | 207 | 202 |
| rRNAs | 3 | 3 | 3 | 3 | 3 | 3 |
| tRNAs | 34 | 34 | 34 | 27 | 27 | 28 |
| tmRNA | 1 | 1 | 1 | - | - | - |

Values in square brackets indicate the corresponding values for the plasmid.

Genome-wide pairwise comparisons revealed extensive conservation of synteny in both symbionts across the three *Diaphorina* lineages. *Profftella* retained highly conserved genome organization in both the Dly–DC and Dly–Dco comparisons, with no evidence for major lineage-specific rearrangements at the scale resolved by these comparisons (Fig. 1a). *Carsonella* showed an almost uninterrupted collinear relationship in both comparisons (Fig. 1b). Nevertheless, several localized differences in gene content were evident among the *Profftella* lineages (Fig. 1a), as detailed below. ANI values were consistently higher between Dly and Dco than between Dly and DC (Table 2). For Dly–Dco, ANI was 94.65% for the *Profftella* chromosome, 90.52% for the *Profftella* plasmid, and 94.52% for the *Carsonella* chromosome. The corresponding Dly–DC values were 90.26%, 86.12%, and 91.90%, respectively. This pattern is consistent with a closer evolutionary relationship between the Dly and Dco lineages, as further supported by the phylogenomic analysis below.

**Fig. 1.**
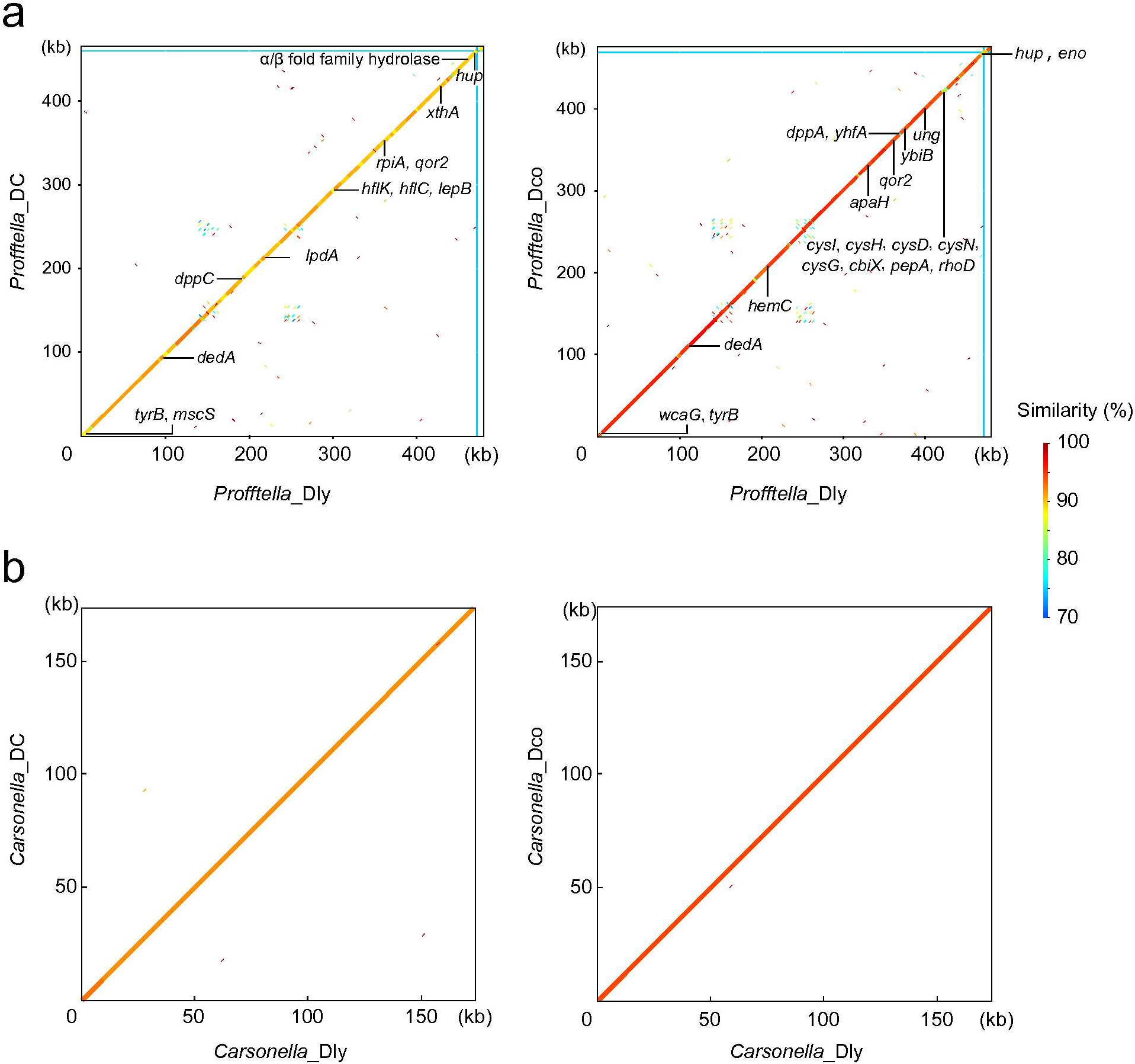
Genome-wide synteny of *Profftella* and *Carsonella* from three *Diaphorin*a species. Pairwise nucleotide comparisons of the *D. lycii* (Dly) lineage with the *D. citri* (DC) and *D*. cf. *continua* (Dco) lineages are shown for (a) *Profftella* and (b) *Carsonella*. For *Profftella*, the chromosome and plasmid were concatenated, with their boundaries indicated by blue lines. The common color scale indicates nucleotide sequence similarity (%). Genes present in one lineage but absent from the other are indicated outside the corresponding plots, on the side of the lineage in which they occur. The α/β-fold family hydrolase lacking an assigned gene name is labeled by its predicted protein function.

**Table 2.** Pairwise ANI values between the *D. lycii*-associated symbiont genomes and their counterparts from *D. citri* and *D*. cf. *continua*.

| ANI (%) | *D. citri* | *D.* cf. *continua* |
|---|---|---|
| *Profftella* chromosome | 90.26 | 94.65 |
| *Profftella* plasmid | 86.12 | 90.52 |
| *Carsonella* chromosome | 91.90 | 94.52 |

### *Carsonella* and *Profftella* show congruent phylogenomic relationships

Phylogenomic analyses based on 26 single-copy orthologous genes independently recovered the same branching pattern for the two symbionts: the *D. lycii*-associated lineage grouped with the *D*. cf. *continua*-associated lineage, whereas the *D*. *citri*-associated lineage diverged earlier (Fig. 2). This topology was strongly supported in both *Profftella* and *Carsonella* and was consistent with the pairwise ANI comparisons. The congruent symbiont relationships provide a phylogenetic framework for interpreting the lineage-specific differences in gene content, substitution patterns, and biosynthetic architecture described below.

**Fig. 2.**
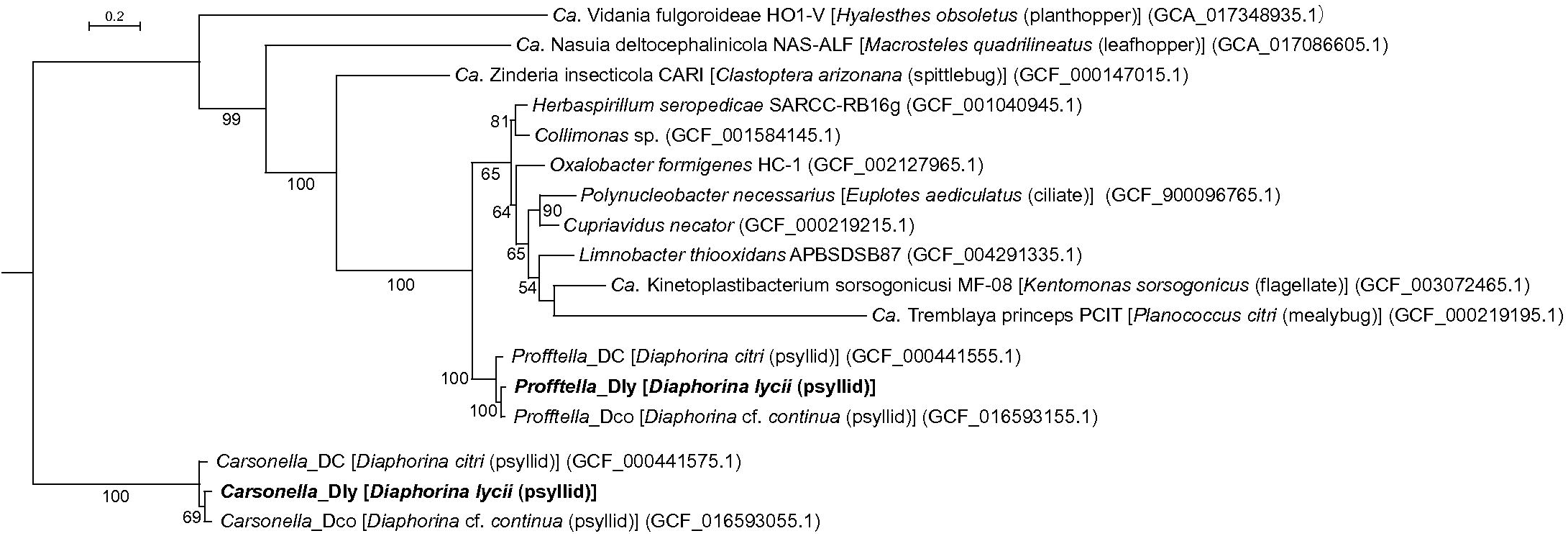
Genome-based phylogeny of *Profftell*a and *Carsonella* associated with *Diaphorina* psyllids and related bacteria. The maximum-likelihood tree was inferred from 4,974 amino-acid residues of 26 single-copy orthologous genes shared among the analyzed genomes. Bootstrap support values >50% are shown. Host species are indicated in brackets for symbiotic bacteria. *Profftella*_Dly and *Carsonella*_Dly, whose genomes were determined in this study, are shown in bold.

### Conserved functional repertoires coexist with lineage-specific gene loss

The three *Profftella* lineages retained conserved gene sets for biotin and riboflavin biosynthesis (Fig. 3). Notably, similar gene sets were retained by the secondary symbiont of *A. mori* and all examined *Psyllophila* lineages [14, 15], indicating functional convergence among independently derived psyllid secondary symbionts. The secondary symbiont of *A. mori* and *Psyllophila* complement the incomplete tryptophan biosynthetic pathways of *Carsonella*. By contrast, all three *Diaphorina*-associated *Carsonella* lineages retain the genes required for tryptophan biosynthesis. Genes involved in the biosynthesis of the other essential amino acids likewise showed identical retention patterns among the three *Diaphorina*-associated *Carsonella* lineages. In addition to these metabolic functions, conservation of the plasmid-encoded carotenoid biosynthesis genes *crtB*, *crtI*, and *crtY* was confirmed in the newly sequenced *Profftella*_Dly genome (Fig. S1; Table S1). These genes are also retained in *Psyllophila*, suggesting another example of functional convergence with *Profftella*, although they are absent from the secondary symbiont of *A. mori* [14, 15]. Phylogenetic analysis of phytoene desaturase (CrtI) recovered the three *Profftella* sequences as a monophyletic group (Fig. S2). A gene encoding a TlyC-family hemolysin, a putative membrane-damaging protein, was likewise conserved in *Profftella*_Dly (Table S1).

**Fig. 3.**
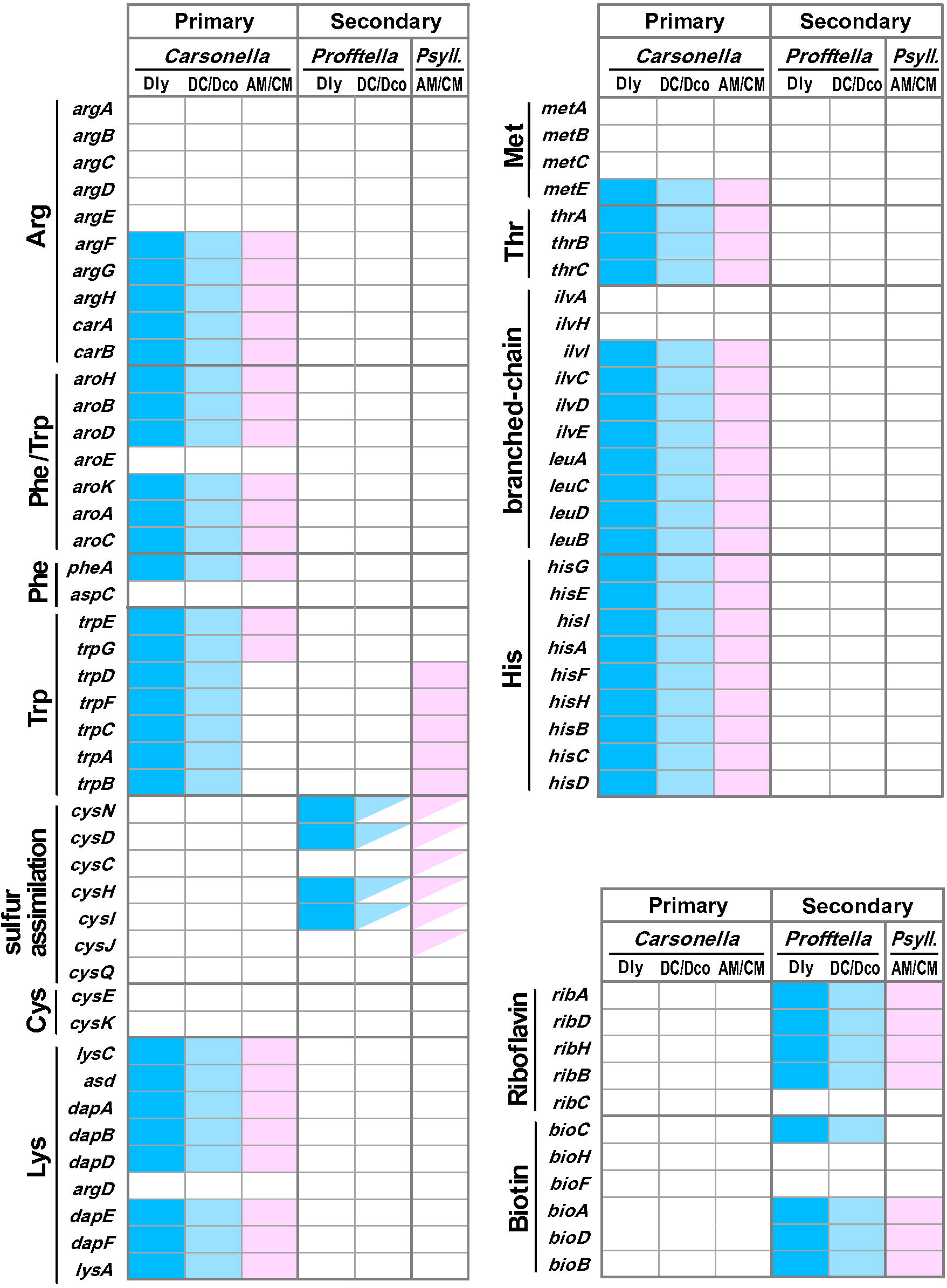
Distribution of nutrient-biosynthetic and related genes in primary and secondary symbionts of five psyllid lineages. The comparison includes *Diaphorina lycii* (Dly), *D. citri* (DC), *D*. cf. *continua* (Dco), *Anomoneura mori* (AM), and *Cacopsylla melanoneura* (CM). *Psyll*. denotes “*Candidatus* Psyllophila symbiotica,” found in many *Cacopsylla* species, and the secondary symbiont of *A. mori*, which belongs to the same monophyletic group. Filled and open boxes indicate gene presence and absence, respectively; split boxes indicate differences in gene retention between the lineages grouped within a single column.

Against this broader pattern of functional conservation, several loci showed lineage-specific loss or degradation in *Profftella* (Fig. 1a; Table S1). *Profftella*_Dly and *Profftella*_DC retained the sulfur-metabolism genes *cysD*, *cysH*, *cysI*, and *cysN*, whereas the entire *cysDHIN* cluster was absent from *Profftella*_Dco (Fig. 3). Sulfur metabolism genes were similarly variable between the secondary symbionts of *A. mori* and the examined *Psyllophila* lineages, being retained in the former but absent from the latter [14, 15]. The *dedA* gene, encoding a DedA-family protein involved in undecaprenyl phosphate recycling, was retained in *Profftella*_Dly but absent or pseudogenized in the other two lineages. Several genes also differed in the extent of fragmentation. The translation elongation factor gene *efp* was fragmented in *Profftella*_Dly and *Profftella*_Dco but remained intact in *Profftella*_DC, although the critical catalytic residue was conserved in all three lineages. The signal peptidase I gene *lepB* showed particularly pronounced variation, being completely absent from *Profftella*_DC, split into two fragments in *Profftella*_Dco, and split into three fragments in *Profftella*_Dly.

### Relative sequence divergence differs between lineage comparisons

We first compared the distributions of synonymous (*dS*) and nonsynonymous (*dN*) substitution rates between *Profftella* and *Carsonella*, using orthologous gene pairs from the Dly–DC and Dly–Dco lineage comparisons (Fig. 4a). In the Dly–DC comparison, *Profftella* had significantly higher *dS* than *Carsonella* (median 0.596, *n* = 361 versus 0.471, *n* = 205; Brunner–Munzel test, *p* = 1.52 × 10^−5^), whereas *dN* was slightly but significantly lower in *Profftella* (median 0.037 versus 0.041; *p* = 0.035). In contrast, in the Dly–Dco comparison, *Profftella* had significantly lower *dS* and *dN* values than *Carsonella* (*dS*: median 0.204, *n* = 355 versus 0.295, *n* = 200, *p* = 3.11 × 10□²²; *dN*: median 0.016 versus 0.025, *p* = 9.64 × 10□¹□). Thus, the relative *dS* and *dN* distributions of the two symbionts changed with the lineage comparison rather than showing a consistent genome-wide ranking.

**Fig. 4.**
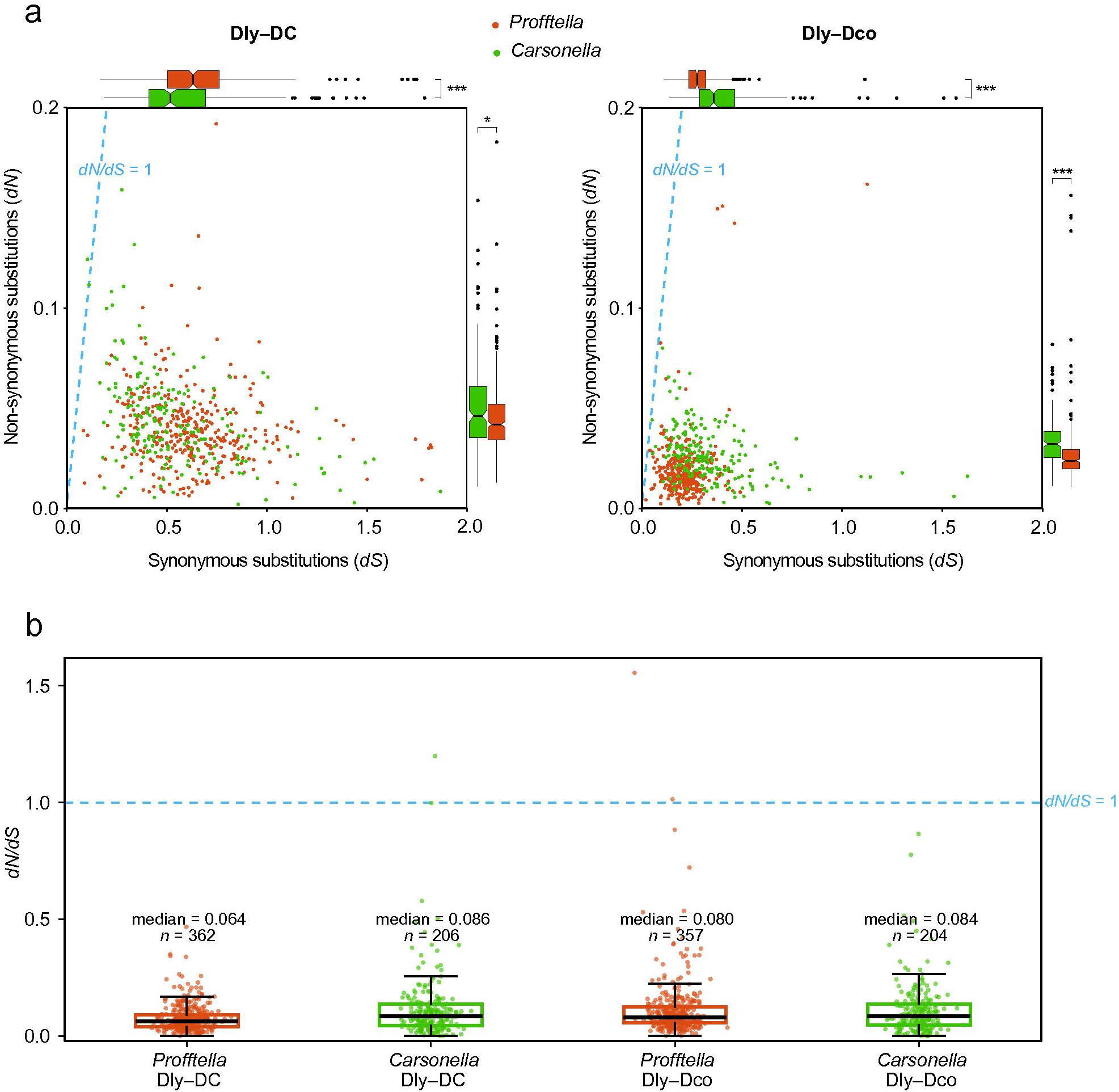
Sequence divergence and *dN*/*dS* distributions in *Profftella* and *Carsonella* across two *Diaphorina* lineage comparisons. (a) Synonymous (*dS*, x-axis) and nonsynonymous (*dN*, y-axis) substitution rates for orthologous gene pairs in the Dly–DC comparison (left) and Dly–Dco comparison (right). Dashed blue lines indicate *dN*/*dS* = 1. Marginal notched boxplots show the distributions of *dS* (top) and *dN* (right); notches represent approximate 95% confidence intervals for the median. Differences in *dN* and *dS* between the two symbionts were assessed by the Brunner–Munzel test (Dly–DC *dN*, *p* = 0.035; all other comparisons, *p* < 0.001). Four outlier pairs (*dS* > 2.0 or *dN* > 0.2) were excluded from the plot for visibility but were retained in the statistical analyses. (b) Distributions of *dN*/*dS* for *Profftella* and *Carsonella* in the Dly–DC and Dly–Dco comparisons. Points represent individual orthologous gene pairs and boxplots summarize the distributions; the dashed blue line indicates *dN*/*dS* = 1. Median values and sample sizes are shown above each distribution.

The *dN*/*dS* distributions provided a complementary view of selective constraint (Fig. 4b). Median *dN*/*dS* values were 0.064 for *Profftella* (*n* = 362) and 0.086 for *Carsonella* (*n* = 206) in Dly–DC, and 0.080 for *Profftella* (*n* = 357) and 0.084 for *Carsonella* (*n* = 204) in Dly–Dco. The great majority of orthologous pairs had *dN*/*dS* < 1 in all four groups. Thus, although the *dN* and *dS* distributions differed between symbionts and lineage comparisons, the *dN*/*dS* distributions were consistently indicative of pervasive purifying selection in both symbionts.

### Diaphorin synthases retain overall architecture but show ACP copy-number variation

The overall organization of the diaphorin biosynthetic system was conserved among the three *Profftella* lineages, but tandem ACP copy number varied conspicuously in DipP and DipT (Fig. 5). In the DipP region corresponding to PedI, which contains two tandem ACP domains, DipP contained three ACPs in *Profftella*_DC, six in *Profftella*_Dco, and five in *Profftella*_Dly. In the DipT region corresponding to the single ACP between the C-methyltransferase and KS domains of PedF, the three *Profftella* lineages contained three, seven, and four ACPs, respectively. A second DipT region corresponding to a single ACP between the ketoreductase and KS domains of PedF contained two ACPs in *Profftella*_DC and *Profftella*_Dco and three in *Profftell*a_Dly. Thus, all three tandem ACP repeat regions showed interlineage copy-number variation, while containing substantially more ACP domains than the corresponding regions of the pederin synthases. The locus-specific KS-domain phylogeny broadly supported inheritance of the *dip* machinery within *Profftella*, with eight of the eleven KS domains showing branching patterns congruent with the genome-scale symbiont phylogeny; the remaining three were poorly resolved (Fig. S3).

**Fig. 5.**
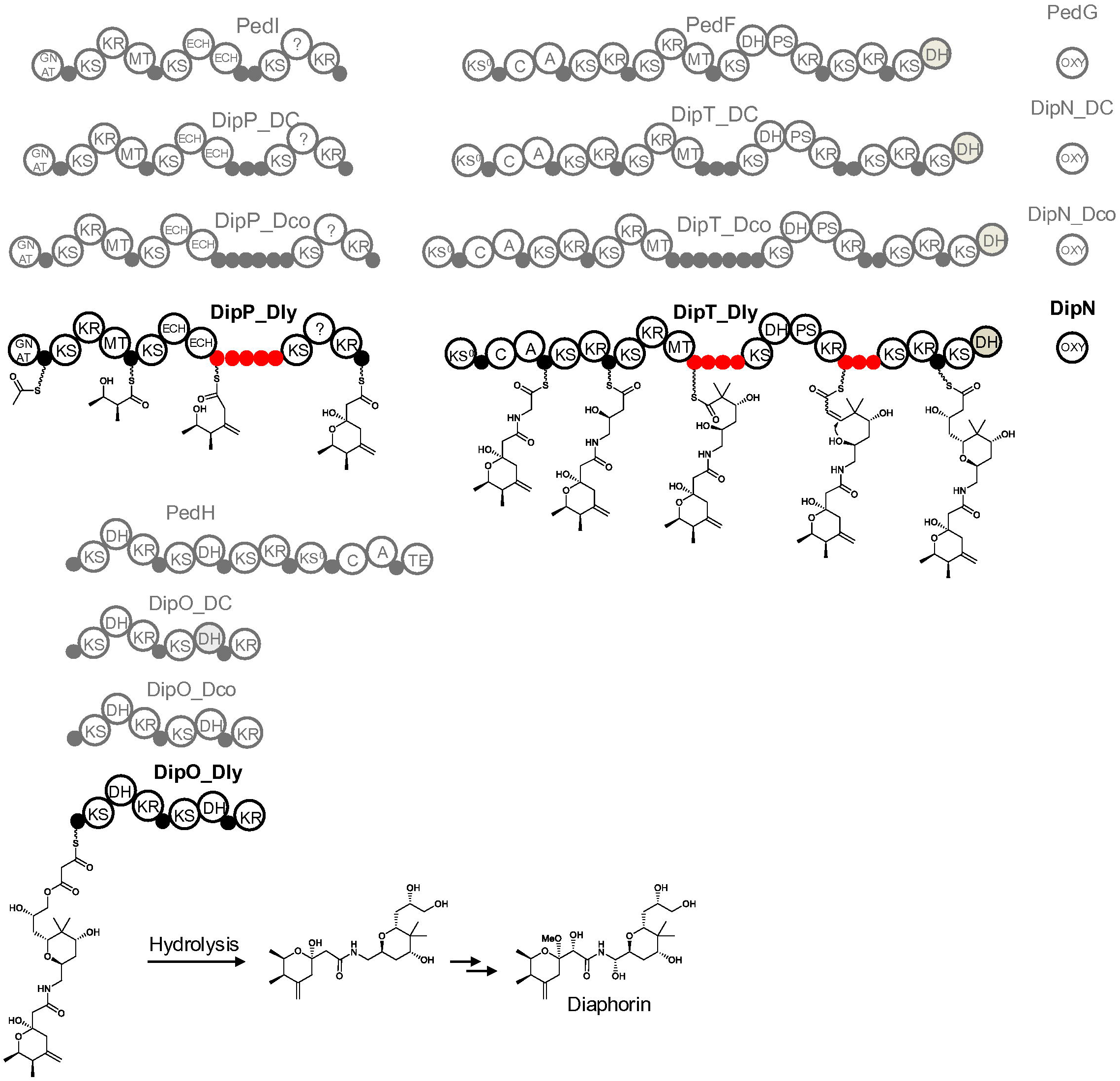
Domain architectures of diaphorin synthases from the three *Profftella* lineages and homologous pederin synthases from the *Paederus* symbiont. The predicted biosynthetic pathway for diaphorin is also shown. Tandem acyl carrier protein (ACP) domains in DipP and DipT from the newly sequenced Dly lineage are shown in red. ACP copy numbers in these tandem arrays are compared among the three *Profftella* lineages and with the corresponding regions of the pederin synthases. Domain abbreviations are as follows: A, nonribosomal peptide synthetase (NRPS) adenylation domain; C, NRPS condensation domain; DH, dehydratase; GNAT, GCN5-related N-acetyltransferase superfamily; ECH, enoyl-CoA reductase-like domain; KR, ketoreductase; KS, ketosynthase; KS0, nonelongating KS; MT, C-methyltransferase; PS, putative pyran synthase; OXY, oxygenase; TE, thioesterase; ?, region of unknown function. Gray domains are predicted to be inactive because they lack conserved active-site residues. Small circles denote carrier protein domains, including ACPs and peptidyl carrier proteins (PCPs).

## Discussion

### Genome-wide conservation provides the background for localized change

Across the three *Diaphorina* species, the highly reduced genomes of both *Profftella* and *Carsonella* show remarkable evolutionary stability, although localized genomic variation is evident in *Profftella*. Genome-wide collinearity is nearly complete in both symbionts (Fig. 1), and their independent phylogenomic analyses place the Dly and Dco lineages as sister groups (Fig. 2), consistent with the higher Dly–Dco ANI values observed for both (Table 2). Thus, despite their distinct evolutionary origins and biological roles, the two co-resident symbionts show congruent phylogenomic relationships and similar patterns of overall nucleotide divergence across the sampled *Diaphorina* lineages.

### Conserved functional repertoires and continuing gene loss

The three *Diaphorina*-associated *Carsonella* lineages retain identical gene repertoires for essential amino-acid biosynthesis, indicating strong conservation of their nutritional capacities (Fig. 3). Notably, the tryptophan biosynthetic pathway is retained in these lineages despite its loss from most other sequenced *Carsonella* genomes [10, 11, 14–16, 18]. In some of these systems, including those of *A. mori* and *C. melanoneura*, the missing functions are encoded by secondary symbionts (Fig. 3) [11, 14–16].

Among secondary symbionts, all three *Profftella* lineages retain conserved sets of genes involved in biotin and riboflavin biosynthesis (Fig. 3). Comparable vitamin-biosynthetic capacities are found in *Psyllophila* [14] and the closely related secondary symbiont of *A. mori* [15], both belonging to Gammaproteobacteria, despite their distant phylogenetic relationship to the betaproteobacterial *Profftella*. Similar capacities are also found in some gammaproteobacterial *Sodalis*-related secondary symbionts of other psyllids [16]. These similarities suggest functional convergence among independently derived symbionts in their contributions to host nutrition. Notably, *Profftella*, *Psyllophila*, and the secondary symbiont of *A. mori* all lack *ribC*, required for the final step of riboflavin biosynthesis. A bacterial *ribC* gene horizontally acquired by an ancestral psyllid may complement this missing function in these symbiotic systems [28, 62]. Thus, *Profftella* retains nutritional functions resembling those of other psyllid secondary symbionts while also fulfilling its distinctive role in diaphorin biosynthesis. The conservation of nutritional gene repertoires in *Profftella* and *Carsonella*, despite ecological differences among their *Diaphorina* hosts, suggests that their core nutritional contributions have remained stable during host diversification. The retention of carotenoid biosynthesis genes and a TlyC-family hemolysin gene across all three *Profftella* lineages further highlights the conservation of genetic capacities extending beyond vitamin and diaphorin biosynthesis. The monophyly of the three *Profftella* CrtI sequences is consistent with the persistence of the carotenoid biosynthetic system through diversification of these lineages (Fig. S2).

Despite the conservation of these diverse functions, several *Profftella* genes show lineage-specific loss or degradation. The *cysDHIN* cluster, involved in assimilatory sulfate reduction, is retained in the Dly and DC lineages but absent from Dco, consistent with loss in the latter lineage. Additional differences in gene content, including those involving *dedA* and *lepB*, are evident in the genome comparisons (Fig. 1a), indicating that gene loss and fragmentation have continued independently among *Profftella* lineages.

### Lower sequence divergence in *Profftella* than in *Carsonella* challenges the primary–secondary symbiont contrast

Comparisons of synonymous (*dS*) and nonsynonymous (*dN*) substitutions revealed contrasting patterns between the two symbionts (Fig. 4a). In the Dly–DC comparison, *Profftella* showed higher *dS* but slightly lower *dN* values than *Carsonella*. More strikingly, in the Dly–Dco comparison, *Profftella* showed lower values of both *dS* and *dN* than *Carsonella*.

More recently acquired symbionts often exhibit higher rates of molecular evolution than long-established primary symbionts. For example, nucleotide substitution rates in several secondary symbiont lineages are one to two orders of magnitude higher than those in their co-resident primary symbiont *Sulcia* [63–65]. Similarly, *Psyllophila* and its close relative in *A. mori* exhibit greater synonymous and nonsynonymous divergence than their co-resident *Carsonella* lineages [15, 16]. In contrast, our previous comparison revealed broadly similar levels of divergence in *Profftella* and *Carsonella* [13]. The present study extends this finding by demonstrating that *Profftella* can exhibit even lower values of both *dS* and *dN* than *Carsonella*. This finding challenges the expectation that secondary symbionts generally evolve faster than their long-established primary counterparts.

Both symbionts exhibited consistently low *dN*/*dS* values (Fig. 4b), indicating pervasive purifying selection on protein-coding sequences. Together with their highly conserved genome architectures, these findings underscore the unusual evolutionary stability of *Profftella* among secondary symbionts.

### Conserved diaphorin biosynthetic architecture harbors striking ACP copy-number variation

The diaphorin biosynthetic system provides a particularly striking example of localized structural diversification within an otherwise highly conserved genome. Although extensive amplification of tandem acyl carrier protein (ACP) domains in *Profftella* was previously recognized through comparison with the related pederin biosynthetic system [13], the three-lineage comparison reveals substantial variation in the extent of amplification among closely related *Profftella* lineages. All three tandem ACP repeat regions examined in DipP and DipT differ in copy number among the lineages, while the overall architectures of these synthases and the organization of the diaphorin biosynthetic system remain highly conserved (Fig. 5). These findings indicate that ACP copy number has continued to diversify within an established biosynthetic framework.

ACP domains carry growing polyketide intermediates during biosynthesis, and tandem ACP repeats have been shown to enhance product yields in polyunsaturated fatty acid synthases and catalytic efficiency in trans-AT polyketide synthases [66, 67]. The lineage-specific differences in ACP copy number may therefore affect the amount of diaphorin produced, although this possibility remains to be tested. Given the experimentally demonstrated bioactivities of diaphorin and its potential role in host defense [12, 31–33, 36], differences in diaphorin production, if confirmed, could influence the defensive contribution of *Profftella* to its *Diaphorina* hosts. Moreover, because diaphorin has been shown to promote the growth of certain bacteria [34, 35, 37], differences in its production could also affect host-associated bacterial communities, potentially in relation to differences in host ecology. Beyond these possible functional implications, the magnitude of ACP copy-number variation is remarkable given the otherwise limited structural divergence of the *Profftella* genomes. The diaphorin biosynthetic system thus illustrates how localized domain amplification can remain evolutionarily dynamic even after extensive genome reduction and long-term stabilization of genome architecture.

### Conclusions

Comparative analysis of three *Diaphorina* lineages revealed remarkable conservation of the highly reduced genomes of *Profftella* and *Carsonella*, including their overall architecture and major nutritional functions, despite ecological differences among their hosts. Notably, *Profftella* exhibited lower synonymous and nonsynonymous sequence divergence than *Carsonella* in one lineage comparison, highlighting its unusual evolutionary stability among secondary symbionts. Against this conserved genomic background, substantial variation in tandem ACP domain copy number was observed within the diaphorin biosynthetic system. This localized diversification raises the possibility of lineage-specific differences in the functional contribution of *Profftella* to its hosts. Together, these findings demonstrate that exceptional genome-wide stability can coexist with continued diversification of a specialized biosynthetic system in an obligate bacterial symbiont.

## Declarations

### Data Availability

The complete genome sequences of *Carsonella* and *Profftella* from *Diaphorina lycii* have been deposited in the DDBJ/ENA/GenBank databases under accession numbers AP054472 (*Carsonella* chromosome), AP054473 (*Profftella* chromosome), and AP054474 (*Profftella* plasmid). The raw sequencing reads have been deposited in the DDBJ Sequence Read Archive (DRA) under accession number DRR1208767 (BioProject accession PRJDB42926).

### Author Contributions

AN conceived and supervised the study. IG and IM collected the insect samples. YY, YH, and AN sequenced and assembled the genomes. YY, JP, and AN performed the bioinformatic analyses. YY and AN wrote the manuscript. All authors read and approved the final manuscript.

## Supplementary material

**Fig. S1.** Circular genome maps of the bacterial endosymbionts of *Diaphorina lycii*. (a) Chromosome of *Profftella*_Dly (471,402 bp; 25.2% G+C). (b) Plasmid of *Profftella*_Dly (8,128 bp; 25.0% G+C). (c) Chromosome of *Carsonella*_Dly (173,860 bp; 17.8% G+C). The blue arcs outside the *Profftella*_Dly chromosome indicate the two diaphorin biosynthetic gene clusters (*dipA*–*R* and *dipS*–*T*). The carotenoid biosynthesis genes (*crtB*, *crtI*, and *crtY*) are labeled on the *Profftella*_Dly plasmid. The concentric rings represent, from the outside inward: (i) genomic position in kilobases, (ii) nucleotide sequence similarity to the corresponding symbiont genome from *D*. cf. *continua*, (iii) nucleotide sequence similarity to the corresponding symbiont genome from *D*. *citri*, (iv) genes on the forward strand, (v) genes on the reverse strand, (vi) dinucleotide bias, (vii) GC skew, and (viii) G+C content. Genes are color-coded according to their functional categories, as indicated in the key. Dinucleotide bias, GC skew, and G+C content were calculated using sliding windows of 100 bp with a step size of 10 bp.

**Fig. S2.** Maximum-likelihood phylogeny of phytoene desaturase (CrtI) from the three *Profftella* lineages and diverse taxa. A total of 515 aligned amino-acid sites from 45 CrtI sequences were analyzed. Bootstrap support values >50% are shown. The scale bar indicates substitutions per site. Source organisms are indicated with higher taxa in brackets. α, β, and γ indicate Alphaproteobacteria, Betaproteobacteria, and Gammaproteobacteria, respectively. DDBJ/EMBL/GenBank accession numbers are provided in parentheses where applicable. *Profftella* sequences are shown in red, with the *Profftella*_Dly sequence determined in this study shown in bold. The *Portiera* sequence is shown in orange, and sequences from other bacteria in black. Archaea, fungi, arthropods, and plants are shown in purple, brown, blue, and green, respectively.

**Fig. S3.** Maximum-likelihood phylogeny of ketoacyl synthase (KS) domains from the diaphorin biosynthetic system and related trans-AT PKS biosynthetic gene clusters. A total of 128 aligned KS domain sequences were analyzed, including the 11 KS domains of DipO, DipP, and DipT from each of the three *Profftella* lineages. Bootstrap support values >50% are shown. The scale bar indicates substitutions per site. Source organisms, protein names (where applicable), and KS numbers based on their order in the biosynthetic pathway are indicated in this order. α, β, and γ indicate Alphaproteobacteria, Betaproteobacteria, and Gammaproteobacteria, respectively. DDBJ/EMBL/GenBank accession numbers are provided in parentheses where applicable. Three cis-AT PKS KS sequences from the *Sorangium cellulosum* epothilone and *Saccharopolyspora erythraea* erythromycin biosynthetic pathways were used as an outgroup. *Profftella* sequences are shown in red, with the *Profftella*_Dly sequences determined in this study shown in bold.

**Table S1.** Gene list for *Profftella*_Dly.

**Table S2.** Gene list for *Carsonella*_Dly.

**Text S1.** Amino-acid sequences and domain architectures of the diaphorin synthases from *Profftella*_Dly.

