## Supplemental Figure 1 for "Dynamic evolution of diaphorin biosynthetic genes against a background of genome conservation in *Profftella*"

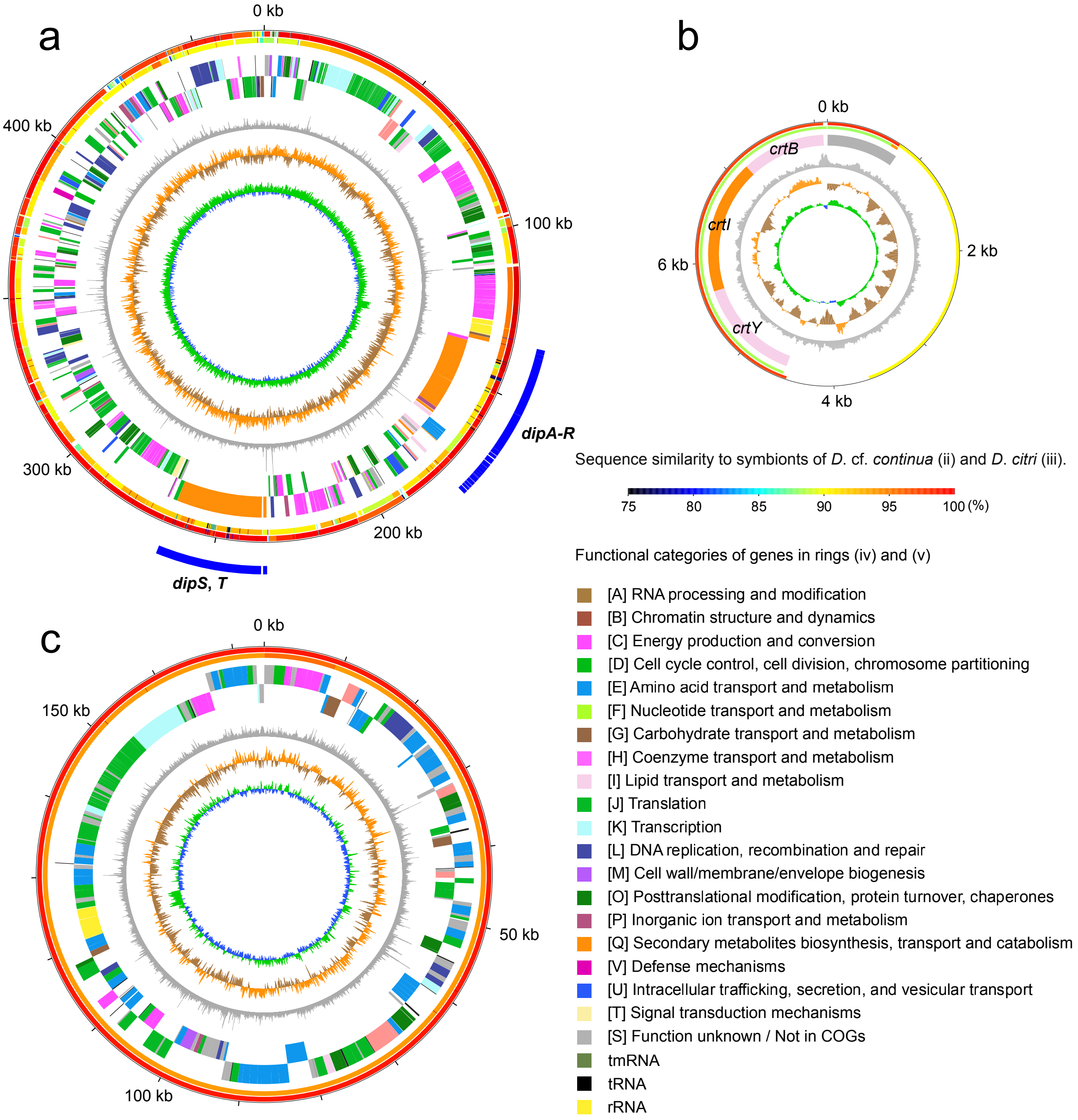
a
0 kb
100 kb
200 kb
300 kb
400 kb
dipA-R
dipS, T
b
0 kb
2 kb
4 kb
6 kb
crtB
crtI
crtY
c
0 kb
50 kb
100 kb
150 kb
Sequence similarity to symbionts of D. cf. continua (ii) and D. citri (iii).
75
80
85
90
95
100 (%)
Functional categories of genes in rings (iv) and (v)
[A] RNA processing and modification
[B] Chromatin structure and dynamics
[C] Energy production and conversion
[D] Cell cycle control, cell division, chromosome partitioning
[E] Amino acid transport and metabolism
[F] Nucleotide transport and metabolism
[G] Carbohydrate transport and metabolism
[H] Coenzyme transport and metabolism
[I] Lipid transport and metabolism
[J] Translation
[K] Transcription
[L] DNA replication, recombination and repair
[M] Cell wall/membrane/envelope biogenesis
[O] Posttranslational modification, protein turnover, chaperones
[P] Inorganic ion transport and metabolism
[Q] Secondary metabolites biosynthesis, transport and catabolism
[V] Defense mechanisms
[U] Intracellular trafficking, secretion, and vesicular transport
[T] Signal transduction mechanisms
[S] Function unknown / Not in COGs
tmRNA
tRNA
rRNA
