## Supplemental Figure 2 for "Dynamic evolution of diaphorin biosynthetic genes against a background of genome conservation in *Profftella*"

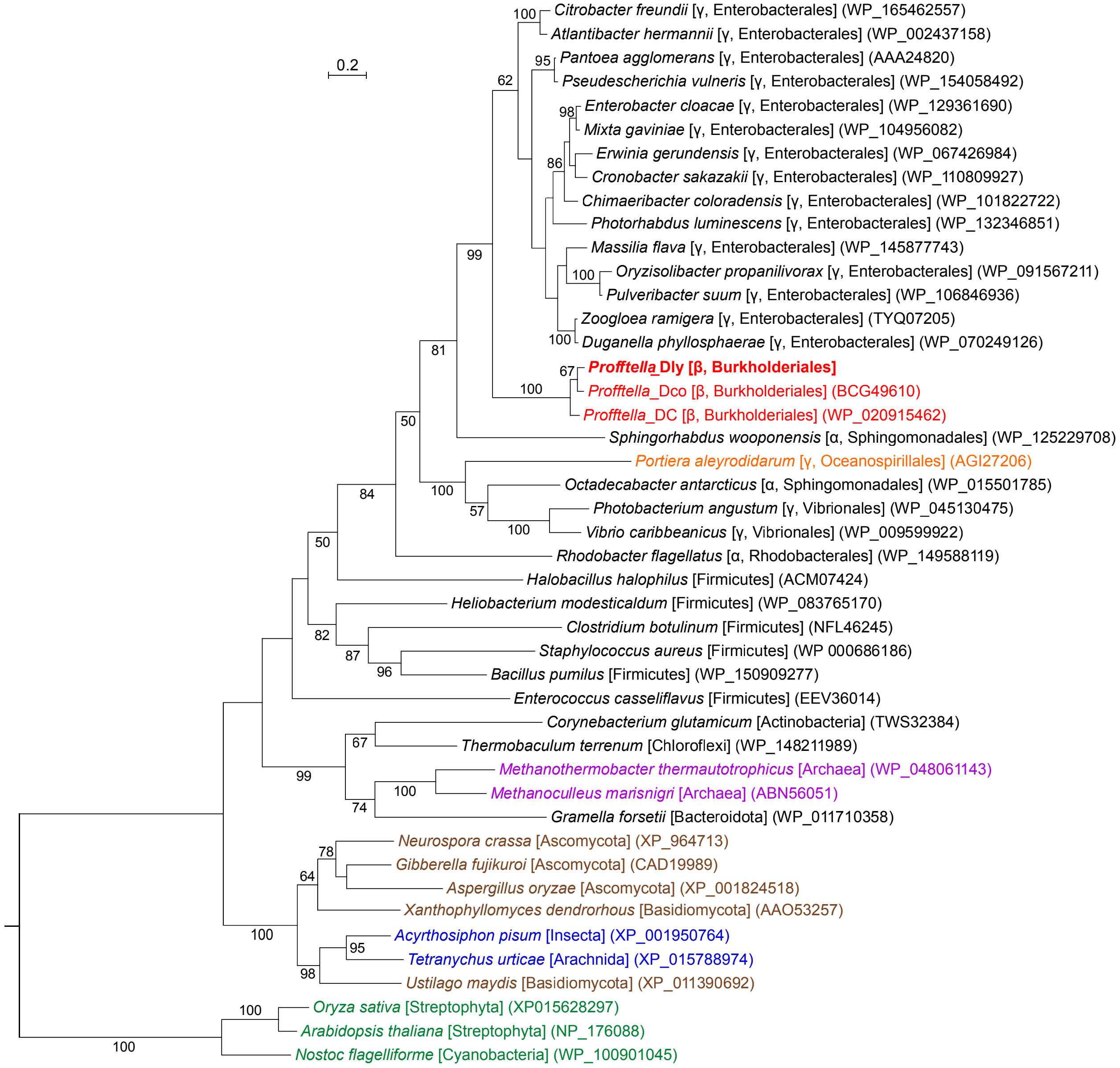

0.2
Citrobacter freundii [γ, Enterobacterales] (WP_165462557)
Atlantibacter hermannii [γ, Enterobacterales] (WP_002437158)
Pantoea agglomerans [γ, Enterobacterales] (AAA24820)
Pseudescherichia vulneris [γ, Enterobacterales] (WP_154058492)
Enterobacter cloacae [γ, Enterobacterales] (WP_129361690)
Mixta gaviniae [γ, Enterobacterales] (WP_104956082)
Erwinia gerundensis [γ, Enterobacterales] (WP_067426984)
Cronobacter sakazakii [γ, Enterobacterales] (WP_110809927)
Chimaeribacter coloradensis [γ, Enterobacterales] (WP_101822722)
Photorhabdus luminescens [γ, Enterobacterales] (WP_132346851)
Massilia flava [γ, Enterobacterales] (WP_145877743)
Oryzisolibacter propanilivorax [γ, Enterobacterales] (WP_091567211)
Pulveribacter suum [γ, Enterobacterales] (WP_106846936)
Zoogloea ramigera [γ, Enterobacterales] (TYQ07205)
Duganella phyllosphaerae [γ, Enterobacterales] (WP_070249126)
Profftella_Dly [β, Burkholderiales]
Profftella_Dco [β, Burkholderiales] (BCG49610)
Profftella_DC [β, Burkholderiales] (WP_020915462)
Sphingorhabdus wooponensis [α, Sphingomonadales] (WP_125229708)
Portiera aleyrodidarum [γ, Oceanospirillales] (AGI27206)
Octadecabacter antarcticus [α, Sphingomonadales] (WP_015501785)
Photobacterium angustum [γ, Vibrionales] (WP_045130475)
Vibrio caribbeanicus [γ, Vibrionales] (WP_009599922)
Rhodobacter flagellatus [α, Rhodobacterales] (WP_149588119)
Halobacillus halophilus [Firmicutes] (ACM07424)
Heliobacterium modesticaldum [Firmicutes] (WP_083765170)
Clostridium botulinum [Firmicutes] (NFL46245)
Staphylococcus aureus [Firmicutes] (WP 000686186)
Bacillus pumilus [Firmicutes] (WP_150909277)
Enterococcus casseliflavus [Firmicutes] (EEV36014)
Corynebacterium glutamicum [Actinobacteria] (TWS32384)
Thermobaculum terrenum [Chloroflexi] (WP_148211989)
Methanothermobacter thermautotrophicus [Archaea] (WP_048061143)
Methanoculleus marisnigri [Archaea] (ABN56051)
Gramella forsetii [Bacteroidota] (WP_011710358)
Neurospora crassa [Ascomycota] (XP_964713)
Gibberella fujikuroi [Ascomycota] (CAD19989)
Aspergillus oryzae [Ascomycota] (XP_001824518)
Xanthophyllomyces dendrorhous [Basidiomycota] (AAO53257)
Acyrthosiphon pisum [Insecta] (XP_001950764)
Tetranychus urticae [Arachnida] (XP_015788974)
Ustilago maydis [Basidiomycota] (XP_011390692)
Oryza sativa [Streptophyta] (XP015628297)
Arabidopsis thaliana [Streptophyta] (NP_176088)
Nostoc flagelliforme [Cyanobacteria] (WP_100901045)
