## Supplemental Figure 3 for "Dynamic evolution of diaphorin biosynthetic genes against a background of genome conservation in *Profftella*"

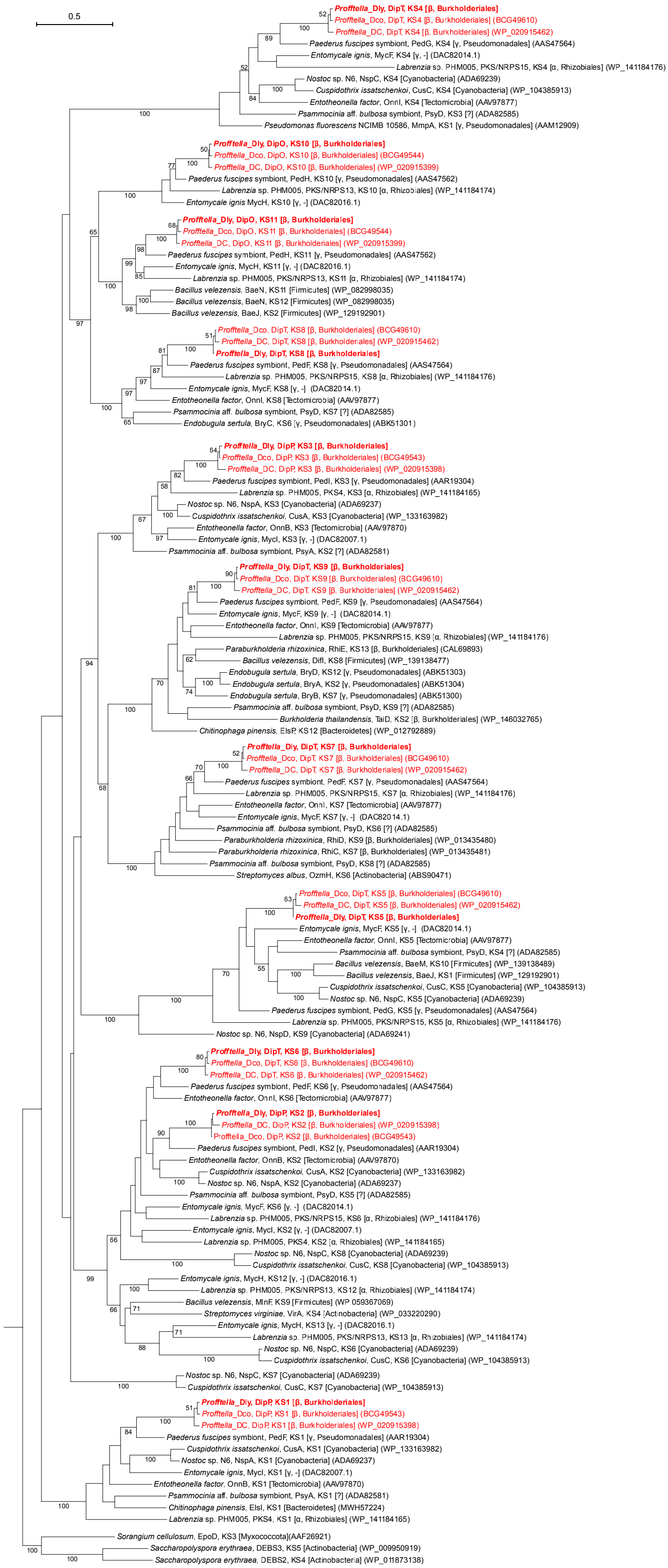

0.5
Profftella_Dly, DipT, KS4 [β, Burkholderiales]
Profftella_Dco, DipT, KS4 [β, Burkholderiales] (BCG49610)
Profftella_DC, DipT, KS4 [β, Burkholderiales] (WP_020915462)
Paederus fuscipes symbiont, PedG, KS4 [γ, Pseudomonadales] (AAS47564)
Entomycale ignis, MycF, KS4 [γ, -] (DAC82014.1)
Labrenzia sp. PHM005, PKS/NRPS15, KS4 [α, Rhizobiales] (WP_141184176)
Nostoc sp. N6, NspC, KS4 [Cyanobacteria] (ADA69239)
Cuspidothrix issatschenkoi, CusC, KS4 [Cyanobacteria] (WP_104385913)
Entotheonella factor, OnnI, KS4 [Tectomicrobia] (AAV97877)
Psammocinia aff. bulbosa symbiont, PsyD, KS3 [?] (ADA82585)
Pseudomonas fluorescens NCIMB 10586, MmpA, KS1 [γ, Pseudomonadales] (AAM12909)
Profftella_Dly, DipO, KS10 [β, Burkholderiales]
Profftella_Dco, DipO, KS10 [β, Burkholderiales] (BCG49544)
Profftella_DC, DipO, KS10 [β, Burkholderiales] (WP_020915399)
Paederus fuscipes symbiont, PedH, KS10 [γ, Pseudomonadales] (AAS47562)
Labrenzia sp. PHM005, PKS/NRPS13, KS10 [α, Rhizobiales] (WP_141184174)
Entomycale ignis MycH, KS10 [γ, -] (DAC82016.1)
Profftella_Dly, DipO, KS11 [β, Burkholderiales]
Profftella_Dco, DipO, KS11 [β, Burkholderiales] (BCG49544)
Profftella_DC, DipO, KS11 [β, Burkholderiales] (WP_020915399)
Paederus fuscipes symbiont, PedH, KS11 [γ, Pseudomonadales] (AAS47562)
Entomycale ignis, MycH, KS11 [γ, -] (DAC82016.1)
Labrenzia sp. PHM005, PKS/NRPS13, KS11 [α, Rhizobiales] (WP_141184174)
Bacillus velezensis, BaeN, KS11 [Firmicutes] (WP_082998035)
Bacillus velezensis, BaeN, KS12 [Firmicutes] (WP_082998035)
Bacillus velezensis, BaeJ, KS2 [Firmicutes] (WP_129192901)
Profftella_Dco, DipT, KS8 [β, Burkholderiales] (BCG49610)
Profftella_DC, DipT, KS8 [β, Burkholderiales] (WP_020915462)
Profftella_Dly, DipT, KS8 [β, Burkholderiales]
Paederus fuscipes symbiont, PedF, KS8 [γ, Pseudomonadales] (AAS47564)
Labrenzia sp. PHM005, PKS/NRPS15, KS8 [α, Rhizobiales] (WP_141184176)
Entomycale ignis, MycF, KS8 [γ, -] (DAC82014.1)
Entotheonella factor, OnnI, KS8 [Tectomicrobia] (AAV97877)
Psammocinia aff. bulbosa symbiont, PsyD, KS7 [?] (ADA82585)
Endobugula sertula, BryC, KS6 [γ, Pseudomonadales] (ABK51301)
Profftella_Dly, DipP, KS3 [β, Burkholderiales]
Profftella_Dco, DipP, KS3 [β, Burkholderiales] (BCG49543)
Profftella_DC, DipP, KS3 [β, Burkholderiales] (WP_020915398)
Paederus fuscipes symbiont, PedI, KS3 [γ, Pseudomonadales] (AAR19304)
Labrenzia sp. PHM005, PKS4, KS3 [α, Rhizobiales] (WP_141184165)
Nostoc sp. N6, NspA, KS3 [Cyanobacteria] (ADA69237)
Cuspidothrix issatschenkoi, CusA, KS3 [Cyanobacteria] (WP_133163982)
Entotheonella factor, OnnB, KS3 [Tectomicrobia] (AAV97870)
Entomycale ignis, MycI, KS3 [γ, -] (DAC82007.1)
Psammocinia aff. bulbosa symbiont, PsyA, KS2 [?] (ADA82581)
Profftella_Dly, DipT, KS9 [β, Burkholderiales]
Profftella_Dco, DipT, KS9 [β, Burkholderiales] (BCG49610)
Profftella_DC, DipT, KS9 [β, Burkholderiales] (WP_020915462)
Paederus fuscipes symbiont, PedF, KS9 [γ, Pseudomonadales] (AAS47564)
Entomycale ignis, MycF, KS9 [γ, -] (DAC82014.1)
Entotheonella factor, OnnI, KS9 [Tectomicrobia] (AAV97877)
Labrenzia sp. PHM005, PKS/NRPS15, KS9 [α, Rhizobiales] (WP_141184176)
Paraburkholderia rhizoxinica, RhiE, KS13 [β, Burkholderiales] (CAL69893)
Bacillus velezensis, DifI, KS8 [Firmicutes] (WP_139138477)
Endobugula sertula, BryD, KS12 [γ, Pseudomonadales] (ABK51303)
Endobugula sertula, BryA, KS2 [γ, Pseudomonadales] (ABK51304)
Endobugula sertula, BryB, KS7 [γ, Pseudomonadales] (ABK51300)
Psammocinia aff. bulbosa symbiont, PsyD, KS9 [?] (ADA82585)
Burkholderia thailandensis, TaiD, KS2 [β, Burkholderiales] (WP_146032765)
Chitinophaga pinensis, ElsP, KS12 [Bacteroidetes] (WP_012792889)
Profftella_Dly, DipT, KS7 [β, Burkholderiales]
Profftella_Dco, DipT, KS7 [β, Burkholderiales] (BCG49610)
Profftella_DC, DipT, KS7 [β, Burkholderiales] (WP_020915462)
Paederus fuscipes symbiont, PedF, KS7 [γ, Pseudomonadales] (AAS47564)
Labrenzia sp. PHM005, PKS/NRPS15, KS7 [α, Rhizobiales] (WP_141184176)
Entotheonella factor, OnnI, KS7 [Tectomicrobia] (AAV97877)
Entomycale ignis, MycF, KS7 [γ, -] (DAC82014.1)
Psammocinia aff. bulbosa symbiont, PsyD, KS6 [?] (ADA82585)
Paraburkholderia rhizoxinica, RhiD, KS9 [β, Burkholderiales] (WP_013435480)
Paraburkholderia rhizoxinica, RhiC, KS7 [β, Burkholderiales] (WP_013435481)
Psammocinia aff. bulbosa symbiont, PsyD, KS8 [?] (ADA82585)
Streptomyces albus, OzmH, KS6 [Actinobacteria] (ABS90471)
Profftella_Dco, DipT, KS5 [β, Burkholderiales] (BCG49610)
Profftella_DC, DipT, KS5 [β, Burkholderiales] (WP_020915462)
Profftella_Dly, DipT, KS5 [β, Burkholderiales]
Entomycale ignis, MycF, KS5 [γ, -] (DAC82014.1)
Entotheonella factor, OnnI, KS5 [Tectomicrobia] (AAV97877)
Psammocinia aff. bulbosa symbiont, PsyD, KS4 [?] (ADA82585)
Bacillus velezensis, BaeM, KS10 [Firmicutes] (WP_139138489)
Bacillus velezensis, BaeJ, KS1 [Firmicutes] (WP_129192901)
Cuspidothrix issatschenkoi, CusC, KS5 [Cyanobacteria] (WP_104385913)
Nostoc sp. N6, NspC, KS5 [Cyanobacteria] (ADA69239)
Paederus fuscipes symbiont, PedG, KS5 [γ, Pseudomonadales] (AAS47564)
Labrenzia sp. PHM005, PKS/NRPS15, KS5 [α, Rhizobiales] (WP_141184176)
Nostoc sp. N6, NspD, KS9 [Cyanobacteria] (ADA69241)
Profftella_Dly, DipT, KS6 [β, Burkholderiales]
Profftella_Dco, DipT, KS6 [β, Burkholderiales] (BCG49610)
Profftella_DC, DipT, KS6 [β, Burkholderiales] (WP_020915462)
Paederus fuscipes symbiont, PedF, KS6 [γ, Pseudomonadales] (AAS47564)
Entotheonella factor, OnnI, KS6 [Tectomicrobia] (AAV97877)
Profftella_Dly, DipP, KS2 [β, Burkholderiales]
Profftella_DC, DipP, KS2 [β, Burkholderiales] (WP_020915398)
Profftella_Dco, DipP, KS2 [β, Burkholderiales] (BCG49543)
Paederus fuscipes symbiont, PedI, KS2 [γ, Pseudomonadales] (AAR19304)
Entotheonella factor, OnnB, KS2 [Tectomicrobia] (AAV97870)
Cuspidothrix issatschenkoi, CusA, KS2 [Cyanobacteria] (WP_133163982)
Nostoc sp. N6, NspA, KS2 [Cyanobacteria] (ADA69237)
Psammocinia aff. bulbosa symbiont, PsyD, KS5 [?] (ADA82585)
Entomycale ignis, MycF, KS6 [γ, -] (DAC82014.1)
Labrenzia sp. PHM005, PKS/NRPS15, KS6 [α, Rhizobiales] (WP_141184176)
Entomycale ignis, MycI, KS2 [γ, -] (DAC82007.1)
Labrenzia sp. PHM005, PKS4, KS2 [α, Rhizobiales] (WP_141184165)
Nostoc sp. N6, NspC, KS8 [Cyanobacteria] (ADA69239)
Cuspidothrix issatschenkoi, CusC, KS8 [Cyanobacteria] (WP_104385913)
Entomycale ignis, MycH, KS12 [γ, -] (DAC82016.1)
Labrenzia sp. PHM005, PKS/NRPS13, KS12 [α, Rhizobiales] (WP_141184174)
Bacillus velezensis, MlnF, KS9 [Firmicutes] (WP 059367069)
Streptomyces virginiae, VirA, KS4 [Actinobacteria] (WP_033220290)
Entomycale ignis, MycH, KS13 [γ, -] (DAC82016.1)
Labrenzia sp. PHM005, PKS/NRPS13, KS13 [α, Rhizobiales] (WP_141184174)
Nostoc sp. N6, NspC, KS6 [Cyanobacteria] (ADA69239)
Cuspidothrix issatschenkoi, CusC, KS6 [Cyanobacteria] (WP_104385913)
Nostoc sp. N6, NspC, KS7 [Cyanobacteria] (ADA69239)
Cuspidothrix issatschenkoi, CusC, KS7 [Cyanobacteria] (WP_104385913)
Profftella_Dly, DipP, KS1 [β, Burkholderiales]
Profftella_Dco, DipP, KS1 [β, Burkholderiales] (BCG49543)
Profftella_DC, DipP, KS1 [β, Burkholderiales] (WP_020915398)
Paederus fuscipes symbiont, PedF, KS1 [γ, Pseudomonadales] (AAR19304)
Cuspidothrix issatschenkoi, CusA, KS1 [Cyanobacteria] (WP_133163982)
Nostoc sp. N6, NspA, KS1 [Cyanobacteria] (ADA69237)
Entomycale ignis, MycI, KS1 [γ, -] (DAC82007.1)
Entotheonella factor, OnnB, KS1 [Tectomicrobia] (AAV97870)
Psammocinia aff. bulbosa symbiont, PsyA, KS1 [?] (ADA82581)
Chitinophaga pinensis, ElsI, KS1 [Bacteroidetes] (MWH57224)
Labrenzia sp. PHM005, PKS4, KS1 [α, Rhizobiales] (WP_141184165)
Sorangium cellulosum, EpoD, KS3 [Myxococcota](AAF26921)
Saccharopolyspora erythraea, DEBS3, KS5 [Actinobacteria] (WP_009950919)
Saccharopolyspora erythraea, DEBS2, KS4 [Actinobacteria] (WP_011873138)
