## Supplemental Text 1 for "Dynamic evolution of diaphorin biosynthetic genes against a background of genome conservation in *Profftella*"

**Supplementary Text S1.** Amino acid sequences and domain architectures of the diaphorin synthases from Profftella_Dly. The domain architecture is shown at the beginning of each protein and corresponds to color-coded parts of the sequence. Abbreviations: A, nonribosomal peptide synthetase adenylation domain; ACP, acyl carrier protein; C, NRPS condensation domain; DH, dehydratase; ECH, enoyl-CoA reductase-like domain; GNAT, GCN5-related N-acetyltransferase superfamily; KR, ketoreductase; KS, ketosynthase; KS0, non-elongating KS; MT, methyltransferase; PCP, NRPS peptidyl carrier protein; PS, pyran synthase; ?, region of unknown function. Domains that are predicted to be nonfunctional (based on the absence of catalytically important residues) are shown in parentheses. In the ACP and PCP sequences, the conserved motif harboring the catalytically important serine residue is underlined.

DipP_Dly

GNAT ACP KS KR MT ACP KS ECH ECH ACP ACP ACP ACP ACP KS ? KR ACP

MHKILENFFKKNENSLEKCFLEAAKIGLFITHQPPKNLLDIKSIKNYILSNLKYKPYCIRPALLEDIPELIKLEKSCWKKLLRTPEFIIKNRICNYPKGQLVLLINNKISGVIYSQLIKSVKELENYNITTVDTLHNIHGSIAQLLAVNIFPDMQQQNLGDQLLEFILIYYSLQAKINLIVAITLCKNFNKLDNLSLSEYIKKRNIYGMLIDPILYFHESHGAIIKKLIPNYRPFDYKNNGYGVLINYDIKNRKCIDINNKIKVKKNNKYNLEEISTYIKRIVYECLSFEFKKFFSIDTPLMDMGLDSSDLLFLNKNLSSHYQLNIDSTFLFYNNTVNKIIFYFYNKINDIDVFKKNNKSYLLGKNFKNIDNNSLNNLNEDIAIIGLACHLPGGINTLSKFWKILKKGSSVIEELHESRWIWPLEIDIKNKHKGINFGAFLKDIEYFDAPFFRISPIEAESMDPQQRILLELSWKTIEDSGYDSNKLSGSNTGVFIGASGSDYTRLLSDYIKIPVNAHHCIGNSMAVLANRISYFYNFTGPSLLIDTACSSSLVAVHAAIQSIKNGESSQALVGGINLILHPFNTIAYYKAGMLSRDGICKTFDKNADGYVRGEGAVMLMLKSLSVALKDNDNIYAIIKGSASTHNGQSNGLTAPNPIQQSYLLQKAWNISNIDARSLGYIECHGTGTKLGDPIEIQGLKNAFHSIFTHEKNIKNFYEKCGLGSVKTNLGHLEAAAGITGILKVILCLKNKKLPASLNFYNLNNNIKLDDSHLYVIDHFQDWVKPRHENGLRIAGISSFGSGGTNAHVVLSEYNIKINNKLINNISINYINSTSPLIFVLSAKTKLQLLIYVKKYIEWLNDKESLNISLRDLTLQLQKGRQAMSCRLALVIFNRNDLLSQLNFFYEKKIIPLQRELDSFNKLKNKRDMVIVINSAIKKKNFKKIAILWQSGIDIDWSLIDKKNIRISLPTYPFSKYYYWLPKNNFNKHISNSDDTHISNSDDTHISNSDDTHISNSDDTHISNSDDTHISNSDDTHISNSDDTHISNSDDTHISNSDVMQLSLTWDPIIMNYENNSYLTNRKLLVIGANFKQKKIIQSINCSMYEFFINITESVDIISYKLNKLISEGYKFDHIIWITPQISKIDFINYQDFGVLLLFKFIKALLSQKIYSSRELIWTIITYATQEVFDTDLVSPGYASIHGLAGVLAKEYPFWIINLLDLEFNTKLEDILVNIPKNITSGETLAYRNGQWFKGYLCSIVNLRLSSKPIYKQKGVYIVIGGSGGIGEAWTNHVIRKYSAKVIWICRSKLDDNIQKKLEKFKKINSSILYVSADACDLTSLKKSLTIIKKIHPVINGVIHSAVSSEFDQSIEKMSIKRFQSILSIKIDASINIAKVFSTEVSDFIFYFSSIVAIEKNGGMSGYSSGGAFEDSYAICLSKKLQCFVRVINWGHWEIGTGEKISNFAKIRLKQSGRLLLKASQAMTFLENISNIPLRQIIVIRVSNPKSIFLFNSLKNLVIYPDNLDIIKPFSSNFMLTESDKKVELLKNLSIFNNKIMEIKLLPLLAGILISTGFLGKDKNNHWKILRNISNNTPKFYEKWLKASIKFLEKYNKKNISIVLENLNNLEKLWESWNISKQKYFNNSNLESAITLVETCLRSLPLILSGKIRSTDILFPNASMRMVNGIYCNNLVSDYFNKILAKTIVSFITLRLQKEPLARIRIIEIGAGTGGTTSIILPYLNPFYKNISEYAYTDISKAFLFHAEKNFLSNYSYITPKIFNIEKPLSDQMIEPGSYDIAIATNVLHATRNICFTLSNVKSVLRKNGLLLLNEISKKSFFAHLTFGLLEGWWLSEDQRIRISDAPGLYSKNWYKILSQEGFNTILFPVKNTHNLGQQIIIAKSNGIIKRKISMEKKIENNQLKSIHKDKNLSLKAIILYLKKIIAKVLHMDISKIDAQELLGNYGIDSILIIQITSKLRKDFSNISSTLLFEYKNIFELANYLIDNHKKELQELTKIKVDNKKVDNKKVDNKKVDNKKYEIKELKINALSKNTIGICDNNDIAVIGMSCHFPKANNLKEYWQLLINGKNCITEIPSSRWEIDKFYNPDPEEAIKQGKTYCKWGGFLNKIMNFDPLFFNISPKEANIIDPQERLFLQTAWETLEDAGYTRERLAKDFNHQMGVFVGVTRTGFDLFGPELWKCGSRLYPHTSFSSFANRVSYFLNINGPSMPIDTMCSSSLTAIHQACQSLRSGESKLAIAGGVNLYLHPSSYILLSSSRMLSKDGICRSFGKNANGFVPGEGVGAVLLKPISKAIEDNDRIQAIIKATHINHGGKTNGYTVPNPNAQFELIKNTLKKANINARNISYVEAHGTGTELGDPIEISSLVKAFRNDTEDNNFCAIGSVKSNIGHLESAAGIAGFIKVILQMKNKRLVPTLHAKIANPNINFSKTPFFLQQKLVNWIREKNIPRVAGISSFGAGGSNAHIIIEEFKKIKKSFHFSCEKKYIIILSARNKESLKNYIKRLYEFTLQMSHEKLANLSYTLQIGREAMDIRIAFIACSLSELQKKLQKCLNIQGNNYSFLEEEENIFYGEIKNYKDVISIFSNDITIQDTLKIWFHKKKYERLLKFWSKGMNIDWTKLYSEKNMYHCIALPTYPFLETPYLLPLFKDKSDFSISNSNQVIDIKYNRLKYINLKDHGKGIFSIKFSDFYNVNENIEEKIKNSCCELISCIKSLSNLDKVKVLILNNLDKLFADTININKLIQTEISNITQSLIKFLFPIIIVLTGKNSNHVMKISMACDLIICSKQGYYRDLSSGIIFNSKSLRKFGLSYPIVDESIIQNYAFEKANQIIRSPSISLKILKKHLNKNIDNIITNLQNFSQYPDFLLKLSYNKDKIFVIPFSSHVIKLEEYYNDILILTICDRENKNMFSSSVVKGIIDAFSYINSSLKYKVIIITGYENYFACGGTKKGLLKIQKGISRFTDEKSYCMPLYCKIPVIAAMQGHAIGAGWAMGLFCDYTVFSEESVYQSPYMCYGFTPGAGSTLIFPQRFGKTLSREILLTAGEFKGKELKERGISMPVLPRRQVLFYSLKLAKKLSLLSRKELELQKKYYSYSLQKALSTVFFHELTMHDYTFVNNPCVITNIERYFDKSTKKNDSKIISNTISYGDSKKLYSSSSVFSILKNSLIEELQIKPEQFDDDAVFIDMGLDSIIAVTWIRKINSQLDLTINSTKIYDYPTLNKFFKFIMSQISEKLQLKNLSINNNIDFDKSTKKNDSKIISNTISYGDSKKLYSSSSVFSILKNSLIEELQIKPEQFDDDAVFIDMGLDSIIAVTWIRKINSQLDLTINSTKIYDYPTLNKFFKFIMSQISEKLQLKNLSINNNIDFDKSTKKNDSKIISNTISYGDSKKLYSSSSVFSILKNSLIEELQIKPEQFDDDAVFIDMGLDSIIAVTWIRKINSQLDLTINSTKIYDYPTLNKFFKFIMSQISEKLQLKNLSINNNIDFDKSTKKNDSKIISNTISYGDSKKLYSSSSVFSILKNSLIEELQIKPEQFDDDAVFIDMGLDSIIAVTWIRKINSQLDLTINSTKIYDYPTLNKFFKFIMSQISEKLQLKNLSINNNIDFDKSTKKNDSKIISNTISYGDSKKLYSSSSVFSILKNSLIEELQIKPEQFDDDAVFIDMGLDSIIAVTWIRKINSQLDLTINSTKIYDYPTLNKFFKFIMSQISEKLQLKNLSINNNIDFDKSTKKKNKKQKNIFLNIFDKNNKKFNNLVNYDIAIIGISGQFPKANNVKKFWNNLLMGRDCVSKIPNIRWFIDDFYDSDKNAIGKTICKKMGFLEDIEIFDPLFFNISPSEAEYMDPQQRLFLQNCWACIEDAGYNPFDLSGSLCGVFVGCAVSDYNRLITNPNKMHNAYALMGESVSILPARISYFLNLQGPCLAIDTACSSSLVAIANACDSLVLGNSKIALAGGVYIINTPDIHIKMSKSGMLSQDGHCYSFDQRANGFVPGEGVGVLMLKKLEDAKKDGDDIYSIIKGWGVNQDGKTNGITAPNTQSQVRLQTEIYKKFDINPEDIQLIEAHGTGTKLGDPIEFEALCESFNKFSYRKNFCALGSVKSNIGHSATAAGVIGIIKIILSLKHKILPPTINYSILNEHIKLENSPFFINTQCKSWNIENKKKRMAAINSFGFSGTNSHIVLSEKIKDNKNIKFINNLPIFTVFPLSAKNLSQLLVYAKLICNYISKDLKNNNINIFDFIYTFQFGRAVMSYRLAVVISSFEELYEKLLNFIKNYKDKENYIDNNFFYGKIKTKKNNTIFKKLKNVERTSENSLEISIILAKRWVDGDIVDWKYLQLFNKNLGYRKHGLPTYPFAREYYWISKKQKKNILYKKNNSKLFNVNYKNNIYNELLDSIDLQDRLQFYKNKQIIFIYTDKNEFNIFKKLLLQLQKVSNLEDKKTNNFISPIDYNFYSFLTEKLFFEEKYEIVLIFNISKTNDDIIIILEKFFQKFLNGYFGCPIEIIFFIQLEISNKEHINKFINYKINLSINNLSKLRRKKIIFCDDYNKFNQLFFIQRLCIEWLISEKNNTLVDFNYIYYLNNKRFIKENLIDKNSKENNIYLISKDWHVKEPYFLKNNIERKILLVLVNEDSIQLVEKILNLNDFKKIILISISNLGIDYISKYDLVNFNDINSARINTQLLVNKYDNNITHIIDLSDIYNVAHDYDNDQFEKIVFYQIIINIINEISILYFTKELQNFKFKKMTLSGSKFSGIIKMISSDYQHINSRCVDIDDSIYNEPLKLRNIIFEEFNIKLQETEICYRDNQRYVPILLEKKLEIKNYMPFLISNKGVYVITGGTNGVGLEIAKYFAEKGCKNMVLMGITNLPPKEDWKKIISNNDLSFYLINKLKELIDLEQKIRNLKIYIGSLDDLNSLEKYFIKIRNNYGPIKGVIHCAGTYSNIKKPGFSNKNIEDIKKVWEPKIKGIENLHKVFKLDPLDFFISFSSMASLIPYLARGSSDYAAANTYMSFFSTYQQFLYKKIFYKNIIISDWNQTGAITRISKEKFIIVEKIFNKLGMRTFNNKEGKILFEKIMNLNNENSTKEDNNSIIINYLNLSIFNNIKERLLYARPFKFKSKLMKKNIEDYIKYWEEKQQNGIKVSIQEIIKIIDINEIKKLSSDFIDRIHKLIIFNKKQKLEKIKENKNFNKQILEKIISKTIIEVLKLKEIDFTETFQSYGLDSISAMVLSTKLEKNLKYTIQPQWLIDFNSVKKLSVYLSTLIVNNK

DipT_Dly

KS^0^ ACP C A PCP KS KR ACP KS KR MT ACP ACP ACP ACP KS DH PS KR ACP ACP ACP KS KR ACP KS (DH)

MKTFQKLRDIETAILQATSHSITNIKENIKPLSIPEPIAIIGLSGMFPKSKSVDVFWESLDSDISLIEEIPESRFNWRHVYNKKFMKENYSKWGGFIPNIDEFDANFFNIPPGEAIIMDPRQRLLLMSVYQTLIDAGYNPEFLKKSNTGVFIAIQENEYLQILKDANIDINEWYSQNCLLANKISYYFDFRGISDIIDAQCPGSAVAIHRAVNALRNKEIEQAIVGSVNILLRPEPFILLSSAKQLSSTNTVNSFGPHADGHIRAEGVSSILLKPLSRAIQDGDLIYALIKNSSVNYNGQGGGSISAPNVESHINLIQDCYQKVNIDPRDINYIEAQGMGNILADLTEWKSFNCALKNIAKQNNIILEDKTCYISTIKPMTGHMESASGLGALFKVIRSMHTKLIHKIIGFKDYHPDMDRENQPCLIANKTIYWLKNKNNSRLAGIHCYGMGGTNSHLLVEEYKREILDTDKDIVPVLIIISAKTKSSLMIMTKKLHNFLAKKNKYRLSDIAFTLQVGREAMKYRISWVVESLQELLVALKKFFESKKFNVLDYTIPVYYGVLEIPYTLQKKLLNIENFQKKKYTKNDLCKIASLWIDGKFYSWNLLYTNKKLYRVRLPGYPFEKQRYWIDNHCKNLCNQEKVSQNIESTTFLIKNKKDNILEIKEYIIKYISDMLQRPQSTIDINQHLYNFGIDSIFTVRLLRNISQKFNIATKGRDLLKNPTINKLSKYFGEILNSSSISIESEKEKRNEHLREYIDYKDSKKQFSLSENQKGLWVLQKIAPNMSAYNVPICFHITRDINLDILQLAFLCTLQKYPILTSIFFQKNKKIIRENKLDKNIIIQKKDVSFIKDKKKIFEYIKTIIKKPFKLDKGPLVRLYLLKGINDYYLLFNIHHIIFDGKSFLPIISFFIKTYSKLLDANKDDTKLIVLNSKINPDDIYEKFVSWESDMLNGVEGNKYRNYWIKQLKNISSTELFTDYPRSTARLFEGEVYSCKLNSSLSYEIKNYAKKQNFNLSSLFLGIFNILISQYTGKNEIIIGMVEMGRSQEIFENSVGYFINMLPIRINNLMYSKESLNKFIDRVQFIIADAIDNSAYPFSLIVRDLRIKPNINFSPIFQIAFEYQNAFSKNDLLEFNNRFRKNFSITMIEEIMQLGEYELVLEIREEVDNFILNFKYNPTLFKISTITRMSEHIVHIIKEIIKNSNLIISEFSILLDKEYKLLEKWNKTIVDYPKELCFHHFFEKQAHISPNSIAVIFKEKKLTYFELNEKSNQLAYYLYDIWKKNSFQKNKLVAVCVNRSLNMTIIFLGISKSGACWLPIDPLYPDERLCFMLNDSKVDIILTEKVLLKRLQNLSNRKKINNSIAIIELDNIQNIFLGNKSLLPKVNSKSLAYIIYTSGSTGTPKGVMIKHRSLTNFLLSMVNKPGFSSKDRLLSVTTYCFDIAYLELFLPLITGGCCCICPSEKLNDAIELKKEIKRLQPTIMQATPSTWIMLFSGKWKNTENLKILCGGEALPLFLNNKFLETNSEAWNMFGPTETTIWSIIARINAKRNENNNFLKKSNNIFIGKPIANTQVLVLDSIGRLSPIGIPGELCIGGDGLAKGYLNNLSLTSQKFITFSFKDKITVYKTGDLVRWSENGMLEYLGRLDQQVKILGHRIELGEIENILNKHIKIKKSVVIARNQQNSKQLIAYYTKKYTEQIPSEELKKYLQKYLPYYMIPSIFINLPDIPLLANGKINRHSLITREIIINQNNSLELEKNIRTNKIINNKENDNILRSKIQYKLLNIWCDLLNIKGIKVHDGFFSIGGNSVLSVILAQKISKDFNIKFHASDLFKYSTIEDISLYIYDCFQKNNESKIINSKISTLKNQSKIRKIVSDDKIVNNFDDYKDCLAIIGISCNVPGAKNYRKFWKNLCEGKESSTKLSLDVLRKIGVPNNILNDKNFIPIQYSIEGKDLFDPEFFNISSKNAMFMDPQFRKLLEHSWGAVEDSGYISNQIPETAVFMSVSNNFYKTLLYNSEMVDNKDEYSSWILNQAGTISTMISYQLGFKGPSVSIHTNCSSTLSGLYLAKQSLQLNEVKYALIGGASLFPIPNIGHTYIPGMNFSSDGHCKAFDFSADGLVGGEGVAVIMVRKAIDAIKSGDHIYAIIRGICINNDGSEKSGFYTPGIKGQTEVITKVLNKTGINPLSISYIEAHGTGTKLGDPAEIIALNDAYKKYTNKKQFCGIGSVKSNIGHLDTASGLIGLIKLALSLKYHKIPPSINYHKPNPEIKFESTAFYVVDRLIEWPGNNIPRAGISSFGIGGTNAHAILEQYKIALENNNFTNFKNTQIIILSAKTNIQLLKIAKNLKNYLEKEIIIDSKYSPQINLENIAFTLQTGRKAMNWRVAFVVDSIEILLNKLKKFISGKDIFENYFYGKIKFEKPIVINNIINNYQDCEYFNKIASDWVSGSLFDWLILHKNKKLNRISLPTYPFKKEHYWVEIPKNEKKVFNSRDYNFLYKPYFYINKNQIIDSKKIIISYPKWCVQNITSLVKNSSNLYFNKYLVFICGLNIENFNIIKSQMIDSQCISLKLNSNDTLIQQYTDVCLKIFNYIKNIISDNINSILIQLILVDRHAQSIFYGVGGLLKTAQLENPNITSQIIDIGDKNTSEFIIKKIIENRYFLKDFFIRYRENREKIIWKIFNDYDKDNLILPWKEGGIYLFSGGSGNLALLFSEEIINKLRSVTLILVYRSKLTLEQEMRITNLRHNKNISIILYQYDVNFLQNVNLVVQDIIKKFKKLDGLIHFSGSIRDNFILKKNTEEFKTVLYDKIAGAFNLDYATRNLNIDFFILFSSIASIVGSVGQADYCAANGFLDAFAYERSIRVKSGKAKGKTLSLNWPLWKDGGMRINKINENIVEKNFGIKSISKNFGLISFYHSLLSKHAQILILEGDKNILEKLLTFKKSNIKDIKKIPKKIFNDHNYDLKQKIIFQLKVLLSKILKTPIEKIQSTELMEKYGVDSIAITQINYELEKIFGNISKTLFYQYQTIDALSNYLISSYSDICSIWGKKFEICNNFLKKSLDIHPNEKSNNIKNFPEKQKSILANNSFFIPLQEPLCIIGISGCYPKSDTLDALWDNIKNGKNCITEIPKDRWSINNFFEKDRKKAVQTGKSYSKWGGFLNGFANFDPLFFNISPKEALGIDPQERLFLQCAWQVLENAGLTREILCKLHFKSLKEKEKFFNQIPRVGVFVGVTKTGFDLYGPEIRKKGKMVFPHTSFSSIANRVSYFLNLSGPSMPIDTMCSSSLTAIHEACQHIRQGDCDIAIAGGVNLYLHPSTYINLCSSYMLSNDGKCRSFGKGGNGFVPGEGVGAVLIKRLSKAIMDRDLIHAIIRSSSINHGGKTNGYTVPNPNAQTELIYMNIEKSGINARAISYVEAHGTGTELGDPIEITGLTEAFRRYTMDIKFCSLGSIKSNLGHLEAAAGIAGLSKIILQMRYKKLAPSLHAREINPNINLETSPFFIQQKLENWTRPVIKINNKNKEYPRIASLSSFGAGGSNAHLLVEEYISDLNKINIEISNTINFPIIILLTAKTKERLEKVVINFLTWLNKKYNLFDNNFLKDIAYTLQIGREHMEERLGIIIKSIEELKNKLNHFLKENVHNSNLGIYRDQINIDKDSFSMLSDDTEFKETIKKWIILRKFDKLLPFWVKGLKIDWEILYLNETYKPKRIQLPLYPFKKEKYWIDSIILKNKKNFNFQENNSQINNSKLIKLLPVWDTINTKELINDFKINYLKKTIILGGNIEQKKEISKYYSRTLELNSILEEDIFTISNKFNNINKFDHIVWIAPVYDINSSHKNNVIIQLFRLIKALLREGYANIDLQITIITVQGLLICDYDEINPDHSSIHSLISSIAQEYPIWSIKAIDMINNNVINWSVSLWYQLLEIKNISIARRKYNWFFQKLIPLNFYISKDIYCKNGVYLVIGGAGGLGCIWSEFMIKNYGAKIIWIGRSPKNNILQEKIDNISIKYNGICPIYIQADARNSDILCSIYHKIKQDYDHINGIVVSTLSDYDLSFKEISEDLFINILSTKRDVSLSIEKCFYKERLDFIVYFSSMVSFEKPGGMSAYVSSCAFSDAFAKKLNKKWNCVVKTINWGYWNIGGGIRISNALKKLIEKRGVESINKEEGMNTLKLFLGNVTAFQQLAITRTNQPEFINIYSNKEFAIVLPNNISNIEKNILNDFHSLENLKAPLKSKDTEELNYWIIRLLFVQLQLLGIFKVNCNYFKKNNAVDIKNQAGILDKYMVWWKEVLNILHEHNYLITSKDTILVNKNISEEVSYTETWRSWKKRKTYFLEKPETHTLAILVNDCLKELPKILHGDILVTDILFPNGSMEKIEGLYKNNCICDYFNDVVSNIVNSYIKNKIINNSMEGIKIIEIGAGTGGTTSTILKKLEPWNNYINEYCYTDISKSFLNHACKFYGSKYPYLVYKLWNIEKSLISQNIDIGDYDIVIATNVLHATKNIRNTIRNVKAALKINGIAILNEISNKTIFASVLFGLIDGWSLAEDTYLRIPGSPGLYSQDWFSLLQQEGYHNISFPIIREHKLGQQIIVAYSNGIIRQDSSLSNQNNSYKLKSNIQNKNFIVEKGIDNIVEKGIDNIVEKGIDNIVEKGIDNIVEKGIDNIVEKGIDNIVEKGIDKSVLINFIQNSILNALSESLMIQINIIKLNIPFSDYGIDSILGVNFIQKINNYLKLKLNTTILFDYTTVDSLTQYIITDYKEIIEKQYLSTVSKEKFVKPIINTNQNSVHKDRMHYSSLSNQNNSYKLKSNIQNKNFIVEKGIDNIIEKGIDKSVLINFIQNSILNALSESLMIQINIIKLNIPFSDYGIDSILGVNFIQKINNYLKLKLNTTILFDYTTVDSLTQYIITDYKEIIEKQYLSTVSKEKFVKPIINTNQNSVHKDRMHYSSLSNQNNSYKLKSNIQNKNFIVEKGIDNIIEKGIDKSVLINFIQNSILNALSESLMIQINIIKLNIPFSDYGIDSILGVNFIQKINNYLKLKLNTTILFDYTTVDSLTQYIITDYKEIIEKQYLSTVSKEKFVKPIINTNQNSVHKDRMHYSSLSNQNNSYKLKSNIQNKNFIVEKGIDNIIEKGIDKSVLINFIQNSILNALSESLMIQINIIKLNIPFSDYGIDSILGVNFIQKINNYLKLKLNTTILFDYTTVDSLTQYIITDYKEIIEKQYLSTVSKEKFVKPIINTNQNSVHKDRIFSKTNFSSSELNLDGIAIIGLSGKFPGAENVNIFWENMISGINPIIELPSHYLDSKYYNIKPQKGKTYCKWGGILSSRDCFDPMFFNISPRDAKSMNPHQRLILQEGWRALEDAGYNPKILSESLTGIFIGAEPTSYIYKTFTGASDAIVASRLSYFLNLRGPSFVVNTGCSSSAVAIHLACQSLRNGETNLALAGGVFAVMNQTILIGLAQTHMLSYSGLCNTFDSGADGMVLSEGVGIVVLKLLKNAIIDKDPIYGVIRASGINQDGASNGITAPSGIAQQQLITSIYKKYKINPERITYIEAHGTGTKLGDPVEANALVNAFKKFTTKTNYCAVGSAKSYIGHTSANAGVTGLISVLLSIKHHQLPGLRNFKKLNPLIQFENSPFYPNIKKSLWKSIDNKPLMAAVNSFGHSGTNAHLVVEEFIFFEKNITNFYKNNSSQLIILSAKDKTRLLKCIKKLFEFITKILSKNNFVKNSENLLSNIAYTLQVGRDSMESRIAFIVNDLNHFIKILKKIIKNNFNINNKFSEDNIWVNEFELVNKITIKDNIIKEWLFCGNLKKLANAWIQGIDIDWEKLYRKNKNYNRIHLPTYSFEKERYWKPNKIKIEKKTLPIFKKNILHPLVHENISTLNKQCFISTFHGHEFFLSDHVINNYKILPGVTYLEMAYFSVNYSINNYSQDKKSFKLKNIIWIRPAFIHNKLLNISINLTKKSDDLINYSIYSDYDISLNKDKLLHNQGLAYIVSSKSESIYLNLSNLKNIFKNFEVEVDKCYMMLEKMNIFYGPSHKSLKIIYCKENQVLAYLCLPDLIKDTYDSYTLHPSLIDGAFQASIALKLNNIDFDNKKTLLPFSLEELEVFGKCEKEMWVWIRHSDNKSLEKNDNKQRLNIDLCTNNGEIRIRIIGFTSRISIDNNIETISYKKNNSISHPLFIKNNFDQTKWTSHFHGNEFFLKDHKNIMPAVIYLEMAYIAASKNTKITGLSNIIWPKILFIKKNNILKTQLIFKNINSIKKYIIFDSQSKGVYCQGNFITDNCDSDQILKNTQVTDFCISDIQLLKSRCYSVLNYQQCNYILQYTHGPSLMSIYKLYSNNKEALALLNFPEKFLDKDNNYVFHPSLLNGAILSSVIWSIINRDKSKLIQNLPMPFSLKFFKIHDISSFSEKLYVYIKKSKNFTLSKNLEFFNIQIFNTKGKLLISLLDLTLIFKEPKYNLLYAVPKWKTEKLSSKIKIINNISKPIFLLIEKNIKLQNALHKVWPNSIIDVIFNKNTQNIEKNIFFEKFLIYIFKYCKNILKENNNIIQSLILLIPLEKKAFLYGGSISGFLKTACFEYTKLATKIIYYPVIKIDDSNQKLSNLIFNISTEISSMSTDVEICFSKEGKRQVRKLCEIDIKNNHENTLFNIKEHDVIWIIGGMGGIGRLLTHYLGIIKKSQLILSGRSELTNKNLKFIEELSSKNINISYLKTNIIEIESINTTLSIIKRKYGKLNGIIHCAGIIEDNYIINKTTSQFMRVARPKIDGLLNIDAATSNLPINYIILFSSIAGFFGNIGQSDYATGNSFMDAFADYRNQCMYSGKRSGKTLSLNWPLWLGGGMQMNVVNEKLMKKATGMTAMDNISGLIAFENVFSKNYNQILIAFGDYSEIKKRLLSFKKDFTYVNFINHEKEIVKFNSNKNLQLTQNEKKLSEMVRSELINIVAKVQHIPHEKISFQKNLSAYGFDSISFTEFANVLNKTYELVLMPTLFFEIPTLIDLESYLFTHHKSELIKKHKSNYIKVKDEKKLSEMVRSELINIVAKVQHIPHEKISFQKNLSAYGFDSISFTEFANVLNKTYELVLMPTLFFEIPTLIDLESYLFTHHKSELIKKHKSNYIKVKDEKKLSEMVRSELINIVAKVQHIPHEKISFQKNLSAYGFDSISFTEFANVLNKTYELVLMPTLFFEIPTLIDLESYLFTHHKSELIKKHKSNYIKVKDEKKILNLKDSLENININTVAIIGIGGKFPGSKDINDFWRKLENNEDAITEVPISRWDWKAIYGDPHLESGKTKVKWGGFLSDADCFDAKFFGISPAEAEVMDPQLRVFIETTWAALEDAGYPPSKLSGSKTAIFAGVSTADYKDILNEARRKGSVKSLAEPFPFMIANRVSYLFNFHGPSEVIDTACSSSLIAINRAIESLHLKNCDLALAGGVNILASPNITIASSKAGLLSENGRCMTFDQRANGYVRSEGVGVLLLKPLKNAIIDNDHIYGIFRGNFENHGGHSSSPTSPNMLAQKQLLVDVYRRSNINPCTISYIEAHGTGTKLGDPIEVNGLKAAFSELHKYYKTPLLRPYCGLGSVKANVGHLEAASGVIGIIKVLLMLKYKKIPGNPHLKIPNSYLKLDNTPFYLVNKTCDWIQLDNNIPRRAGISSFGVGGSNVHVIIEEYIQNIKINGIKENNILIPIIILSAKTKESLKRYVMLLLEFIIKEKNNFSLYDLAYTLQVGREAMNYRLAIHVNSYEDLIKKLQDYLNKKITHGIYINYSEKDKNENLEKINKSKNISILIDSWVKKNRYDKLILSWINGSEFSWDKLYNIFKPKRINLPTYPFDRQKYWVSIKQKLVNQYDNTPIINEEEKIYDKKSIFTFEEVWVKSPLIFLPFSKVKKKNLLCFSSNKDFQSKIIKSVSNYTNIIFIEQINNIKDIQGPLGHTVNYQINKDNINDYISFLNKIYKKDKKIDFVLYLWPLEDKNCIYNTMPIIYFLQAVKKSKLYLHKLIISGISINKKNEENLYYCYFNSWIAFERSLHIIMPETQISIVFAEKITNFNIQSWITILWKELQEPKLFSIMYYGNDLVRYTLNIKSIILKDFKNSYIKQGGTYLITGGLGRLGMLFSNYLINKYSTKLILSGRSKLDSIIHNKLKKFNGKAIYIQVDVSDKFKMISEINSIINNTGPIDGVLHIAGISGLTSNILEANYKNFYSVLNSKISGTIALNYALSNTICKLQNNKLDFVCYFSSSSAILGDFGSCDYAMGNRFQTAYAKYLNNQNIYKKYTNKLIVINWPLWNNSRFKIGNHEQADFYLKSSKQKILYENDGLKLFEKLLIQDKTQYLVLVGELEKLNQIINNISPKISENKKINNISPKISENKKINNISPKISENKKINNINIEKFIENDLKNHICNILNTKNNEIYKNKNLADYGFDSISLAEFSRILSKFYSLDIMPSIFFSYSTLERLITYFIKNHNDTMIEFYRNKYSFKIVDKKNISTLYADKINKNFKNISKKSNFILNKNIDIDSEDLIAIIGMSGRFPSARNINDFWKILINNKDVIEEIPRKIFDWRLYYENPIKSSNKINSKWYGSIPGIDEFDPLFFEISPLEAERMDPRQRHLLQESWLALEDAGYGPNQISNQKIGMFVGVEEGSNYQDRLDQVNLTSTHNAILSARLAYFLDLKGPVMAINTACSSSLVATHVACQSLRQYECDTAISAGVNLMISPEAYIAMTNAGMLSPNGKCYVFDERANGLVPGEAVVAIVLKRLSHALSDGDPIHAIIRGSGINYDGKTNGITAPNGISQTELIKSVYKKSNINPEDINYIVTHGTGTKLGDPVEINALYDVFKNKTCNQNFCAITSNKSNIGHTFAASGLVSLINLVQSIKYKIIPASLHCEKENNYIVWKNSPFYINKSNKKWDVVNEKLRIGAVSAFGMSGTNAHIVLQEYISTNFKNIKNNIFISSNPYHIVVLSAKTKISLKEKMKQILLFLKKNNTVCIENFVYTLMQGRYHFQYRCAIIISSIKEIIKIINDILSKEKILYTNYLQGIVSHKFTSNKIICSYINELTKRCLLSKIQKKEYYENLLALGDLYCQGYIIPWNNLYPNKFERIHLPGYLFLKEHYWIEKENINSLKEKNLKEKNTEYRRIEFSIQLLNLSRIILEKKINNSINILKNIIWGKLPEIENYKDIKLKINVYKKNNKLVYTTNIVKHCCQIGEIDLNNTIFIKNPIDFNKLQKDLYKLKINCKNVSKIYISKQFLFADILLNKKNNSYSESIFFSKALELIHVFVDYHKIIINNDLLPFSLKYIKIYKKIPLNVILYLICKKVEPNGSYQQYDIIFYDSKGNICLKLKDLIFKKSNKLINIDEIFSNL

DipO_Dly

ACP KS DH KR ACP KS (DH) ACP KR

MIFSNFQDINIKMKKFLENEIINSSKSDLENNFKNSKKKLVTKNKEKLILNIEKKEKNSLYICFYKKIANIVSKILKIPLERMNMQENISRYGVDSIIVTEIIKCISDILNFPIAPTVFFEAKNIEELVNILFQRYKEKIKLYFLKNHKRNQALEAKVNQEPKKQETKENNTEIKDWINKFKFIIASSDSINASSDSINASSDSINASSDSINASSDSINASSDSINASSDSINYEPIAIISMEGIFPNSPNLQTLEKNLRNESDCISEIPLSRWDWKKIFGDPQTGNFTNVKYGGFAPDIDKFDPIFFGMSPREAELMDPQHRLFIQCVWKLIESAGYAPKSLSGKKIGIFIGINLQDYAHIVNKSRKMDSLHLTSLGHMFCPNRLSFYLDIHGPSQVIDTACSSSLVALHRAILSIQHEGCEMSIAGGANLIISPDMHIMYNKIGMISKDGRCKTFSKYADGYGRSDGIGVVLLKSLRLAEKDNDNILAVICGSSENHGGMSTSLTAPNPKSQAKLIIEAHNRAKSDPRSIGYIECHGTGTKLGDPIEVNGLKMAFEELFRINNLSKSNIKFCGLGSIKSNIGHTETSAGIAGVIKTVLSLRNNYLYKSLHSEDINPMIELEESPFYILQKGCLWKSPILNNNKLPRRAGVSSFGAGGSNAHIVIEEYIKHTKVIEKKCNFKVIDSDVLIILSAKNIDRLNDIIIEMFYFVKKILAKKKVVNLVDIAYTLQVGREAMSERLAIIVNSISQLYLKLEQIYFSIKNKEKLGNIISCYYGNLKKEKNIFYVKNEHFISDNDINNCFNNKNYKQLALWWIKGVTINWSKLYANKKSKLHRVSLPTYVFSKQSYWIENLDNKKNIDYYLKSHNLLKNSISNLGKYYFSAKLTGKEFFLADHIVMNKKILPGVIYLEMVCSAIKEIKIKNISNQFIIQLKNVVWAQSFFINKNCSKVLHMKMNQETNDQISYQIYSEQEKNNINVRKERIIHSQGIAITKNIPIINDINDINIDNLLENFTKRQLDTNFLLGDICYQMFNDMGINYGPTHQCLEKIYFSQKNEKNSPQVLAKLKLLKNIKENYKYSKKFMFHPGLVDSGLQACIGFMAYFGYINFDKKTNKVIINNKNIVSLPFALEYFLLLSQPSTSMWVWIRYSKNVLFNSNIQKLDIDFYDKQGKVCICMRGLSSRSPKTNNLEKDFTTLVYKPIWVREDIKIGIKKNKIKNLSNFSIWSHHFIILCDFNNNFPKNIEEDIKKIIPFSKYFYLCIQGSIQDIYVTISLQIFRKIRKILRSKLQGNILLQIVFIENEKSMFLNGLSGLLQSVSRENTKFYSQLIGLRDYNTLNLYSSEYITNIIYSNSVGKVYKNKIRYSSIGREVFSWKELNFISSTKAIFSPLKENGIYLITGGNGGLGLLIAEFITQKINKGVVVLCGRSFLNDKIQKNINNLSTKNIKIKYYCLDITKKIEVEEFIKNIIHKYNFINGIIHCAGLLMDNFVYKKSSSEFIKVLEPKVNGTIYLDEATKTLNLDFFILFSSISGSLGSAGQSDYATANSFLDIFSDYRNKLTSRNQRYGRTISINWPLWSNGGMKMQESVKNIMLQTSGIVEMPNINGLDILYKVITLNISRIMVIYGISSRIRNYLENIDKKVINSLFQENKSSSIIDSDNINYNELRNRICKMLSKYISKLMKFSLEDIENDVQLSDYGFDSITFTDLANRLNRKYQIELTPATFFEYPTINALSELLSVQYYDKFIFKFNLKPKFIEKFEKLDKELYLSTDSIEKKEKLLPISDNKLKNKSHSLLDNKIAIIGISGCFPMSRDIDEFWKNLLSGADCISEIPLSRWDWRAIYGDPKKDANKTNIKWGGFIDNIDVFDASFFNISPREAELMDPQHRLLMQYVWKAIEDSGYSPGSLSGSKTALFIGTASSGYGQLIANSNFTIDSYSATGVIDSIGPNRMSYFLNLHGPSEPVETACSSSLVAIHRALSAILMGDCEQAIVGGINLIISPETQISFSKAGMLCEDGRCKTFSNKANGYVRGEGVGMLFLKKLSAAEKDGDHIYGVICGSSENHGGRSSSLTAPNPKAQTEVIKDAYIRANINPQSISYIEAHGTGTELGDPIEVNALKSAFRDLNKINNNSEKFIKNAYCGIGSVKTNIGHLELAAGIAGVIKVLLQMKYKTLVKSLHCEKINPYITFKNSPFYLLHNNCKWKRLRDINGKKIPRRAGISSFGFGGVNAHLVVEEYSIKKEKFKYESNSIALIVLSAKNIESLKEYAISLIKFITINNEISIENKIYDHDNICKFLSNIVRKYISDILNINKKEINFDDQLSQIGLDCIYGVMLLEKLSKKFGIDISFNVLLNDHSIISFVNAILLNYPNLYAIIDKKYKKKQILNFDNHFKKIRNDINLFELAYTLQVGRDPMDHRLALTAISFDELLIKLQAFIDDRINDFQGLYLGRIKENKRIFSIFINDKEIQEALEKWMKCGKLTKLLEIWVSGMKISWEKLYGESCLYPKKPKRISIPGYPFSKKSYWIKKIFKKKKIYDFDSRIVEKLKKLHPLVHENISTLKYCKFRTFLTGEEFFLSDYSINEQKILPNIVYLEMVRAAAVIAQDNKHSDVIEFHDIIWTNPFVFSEEKYIEIELNKNIGNDIQFKIISYSTLDNINFSKLQIHSQGVISYNNSNKHNLPCVDINLILNESNIQKFSSVQCYNLFKKFKLIYGPTYRNVHELYVSKKQVLAKLILPELMLVTLKKYILHPSIIGSALQCIIGLYLDKLGLTSTEENNKELFLSFVIEDIYIFDICTKLMWVHIEYASSHLTKEHIKKININLCNEHGVIHIQIKGYILSKFNFENKEYKINFNNKKLNSKNINNSIELILDKSDKNIDKLLYNKTIKYFKEIFSSLLKYPIDELDINENINSYGINSMAIIELNVILEKQFGTLSKTLFFEFNTLNSLTNYFISSYHSQLIKLFNINNIKKNKKEKSLSLCQSKIIPKKKNTKIKISSLVKKYSILITTPKWKESSISSLPLNKNNKNNIVNKKNYDTHVIFFTNEIILIPGAVCLSLFSIIKKIDYYFFDCSIKIFEYIQQNLINKNGRILLQIVIQNFGSKICSNITFPVKFLHSLSGILKTAHIENSNFFGQIIEITHQKNKDNFIQEIIQANALIPFDQNICYLNGKRYVSYWDKISSVKNIFPFFWKNDGIYLITGGMGKLGLIFAKEIINQTNNVTLILIGRSKLDSLKQKKLNSLKIRGAEIEYYSIDVTNQTAILKLIKSINLKKNHGLNGILHCAGIIKNNLIYKKSIIEFKEVLLVRVLGAINLDIASRGMNLDFMVLFSSISGVFGNYECCDYSTSNFLVNMYSDYRNQLSLSSFIASQNPRGRTLSINFPLCNNNKINIDNYNKELVYKDIDIKKLNTQNILSIFYYAMHTSYSQIVILQGNLPYLRKLLFLNKKTPQKKNSKKLSDKKLSDKKTPQKKNSKKLSDKKLSDKKTPQKKNSKKLSDKKLSDKKTPQKKNSKKLSDKKLSDKKTPQKKNSKKLSDKKLSDKKTPQKKNSKKLSDKKLSDKKTPQKKNSKKL
